# Molecular simulation of ion-mediated phase separation and structural organization in RNA condensates

**DOI:** 10.64898/2026.11.13.766555

**Authors:** Lipika Baidya, Heyang Zhang, Hung T. Nguyen

## Abstract

Phase separation of RNA is strongly influenced by ions, yet the molecular mechanisms by which ion-RNA interactions affect RNA condensation and internal organization remain poorly understood. Here, we use coarse-grained molecular simulations to investigate how temperature, ions and RNA sequence collectively regulate condensate formation and organization. We find that RNA condensates selectively enrich Mg^2+^ through dynamic multidentate phosphate coordination, producing ion concentrations several orders of magnitude higher than in surrounding solution. Rather than creating a uniform dense phase, this ion enrichment establishes spatial gradients in ion-RNA interactions that generate distinct microenvironments in the condensate. RNAs in the condensate core accumulate more Mg^2+^, exhibit enhanced base stacking and transient helicity. These heterogeneous ionic environments emerge spontaneously even in condensates composed of a single RNA species and are further modulated by RNA sequence through differences in Mg^2+^ binding and packing in condensate. Together, our simulations provide a molecular-level framework for understanding the origins of heterogeneous organization in ion-mediated RNA condensates.

**For Table of Contents Only:** 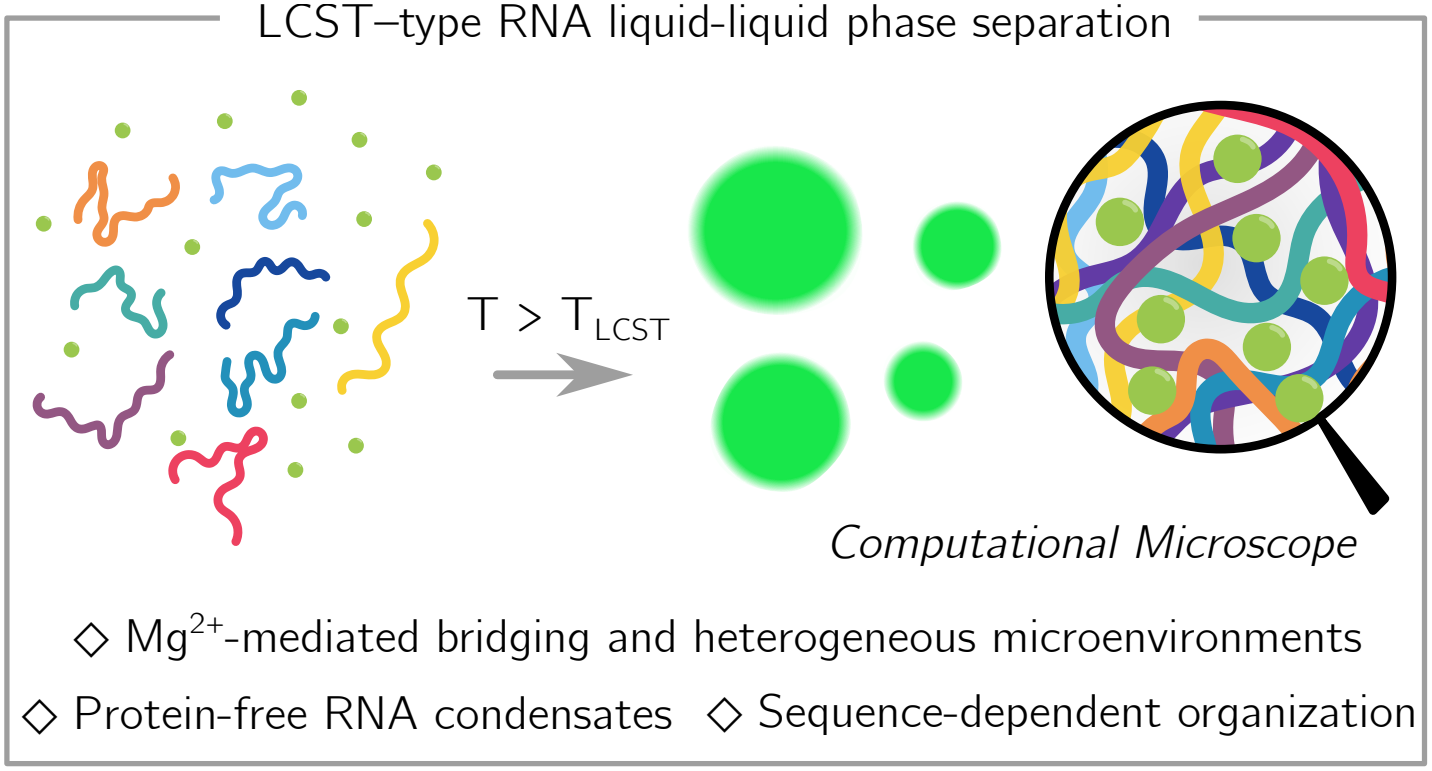

## Introduction

Biomolecular condensates formed via phase separation (PS) have emerged as a fundamental mechanism for intracellular organization, enabling the spatiotemporal regulation of biochemical processes without membrane confinement.^1–4^ PS gives rise to diverse cellular assemblies, including nucleoli,^5,6^ stress granules,^7–9^ nuclear speckles,^10,11^ processing bodies,^12,13^ where biomolecules (e.g. RNA, DNA, protein, etc.) function both as scaffolds and regulators of condensate material properties,^14–16^ dynamics,^17,18^ and function.^19–22^ Dysregulated PS can potentially lead to solid-like pathological aggregates associated with neurodegenerative diseases. ^3,17,23–28^ Notably, many of these cellular condensates exhibit multilayered organization, with distinct sub-compartments that selectively recruit specific proteins and nucleic acids to carry out specialized functions.^2,5,29–31^ Increasing evidence indicates that condensates are not compositionally uniform but instead contain distinct physicochemical microenvironments that influence client recruitment,^29,32,33^ reaction kinetics,^22,34–37^ and condensate aging.^38–41^ These microenvironments have been shown to exhibition-specific effects, and the Hofmeister series framework has proven useful for rationalizing such behavior.^42–48^

Among the macromolecules that drive PS, RNA plays a central role owing to its polyelectrolyte nature, structural versatility,^49,50^ and capacity for multivalent interactions.^18,51^ Previous studies^32,33,52–55^ have investigated the physicochemical microenvironment of protein-rich and protein– RNA condensates, providing important insights into how spatial heterogeneity regulates condensate behavior. RNA itself can form condensates through homotypic (RNA-RNA) interactions in the presence of multivalent ions such as Mg^2+^, Ca^2+^, spermine and spermidine by screening backbone electrostatic repulsion and mediating specific ion-RNA interactions. ^51,56–62^ Ion-mediated RNA PS is hypothesized to be driven by dehydration of ion solvation shells upon RNA association, a process that becomes more favorable at higher temperatures and can therefore give rise to lower critical solution temperature (LCST) behavior.^51,63,64^ Experimental studies have reported both LCST and UCST (upper critical solution temperature) behaviors in ion-driven RNA condensates, depending on RNA sequence, chain length, ion identity and concentration.^51,56,58,60,65^

Despite these advances, limited spatio-temporal resolution of current experimental techniques is unable to probe the microscopic origin and resolve RNA structure, ion distributions, and their internal organization within condensates. Recent coarse-grained simulations suggested a strong favorable entropic contribution in RNA PS.^66,67^ On the other hand, atomistic simulations are largely restricted to very short RNA sequences (e.g., tetra-nucleotides) owning to high computational cost and limited accessible timescales.^68,69^ As a result, they are limited in capturing how temperature and ions jointly regulate ion binding, RNA-ion interactions, ion partitioning, and ultimately drive PS. Several key questions, therefore, remain open: (i) how do temperature and ions together regulate RNA structure and ion-RNA interactions in the condensate; and (ii) how does the spatial localization of RNA and ions define the microenvironment within condensates?

We have recently extended our previous framework for RNA simulation^70,71^ to include explicit ions and temperature-dependent ion-RNA interactions,^67^ which quantitatively captures RNA structural properties, ion binding, and temperature-dependent ion-induced collapse of RNA. In this work, we report the application of this model for the investigation of temperature- and ion-driven RNA PS. Our approach captures ion solvation and distinct modes of ion-RNA binding, allowing a systematic examination of how temperature and ion conditions influence RNA structure, ion partitioning and binding modes, while also resolving spatial heterogeneity within condensates. To decouple ion-RNA interactions from RNA structural (un)folding, we consider disordered RNA sequences (polyadenylic acid rA_30_ and polyuridylic acid rU_30_) as model systems, which have been widely studied both experimentally and computationally.^16,53,55,66,72–74^ Our simulations reveal that RNA adopts different conformations within condensates, surprisingly gaining transient structures compared to those in the dilute phase. Condensates accumulate a higher concentration of ions, which stabilizes more compact and helical RNA conformations due to enhanced base stacking interactions. Strikingly, despite containing a single type of biomolecules, we observe internal heterogeneity emerging in RNA condensates:^41^ RNAs in the core bind to more Mg^2+^ and thus are more helical than those near the interface. These core-shell differences arise from spatial variations in ion-RNA interactions, which can be tuned by temperature, RNA sequence, metabolite and ion concentrations. Our findings align with previous works^36,42,75–78^ which demonstrate that condensates constitute a distinct physicochemical environment capable of modulating RNA structure and potentially affecting biochemical processes that require specific local conditions. Overall, this work highlights how the coupled interplay of temperature, ions, and RNA sequence can govern emergent mesoscale organization, paving the way toward predictive control of biomolecular condensates for designing functional synthetic compartments and understanding their roles in health and disease.

## Results

### Ion-mediated, temperature-controlled phase separation of RNA

Our model has successfully reproduced the structural ensemble of rA_30_ and rU_30_ at varying concentrations of Mg^2+^, in quantitative agreement with small angle X-ray scattering (SAXS) profiles.^67^ Using the exact model, we observe that rA_30_ undergoes an LCST transition from a dispersed solution to a condensed droplet state upon heating in the presence of Mg^2+^ (Fig. 1a-b), consistent with recent experiments^51,56^ and simulations.^66,68^ Representative snapshots illustrate a dilute phase comprised of dispersed chains and a dense phase containing RNA-rich condensates, with Mg^2+^ ions present in both phases (Fig. 1a). The phase diagram (Figs. 1b, S2) shows that increasing the temperature facilitates PS, and high Mg^2+^ concentrations further stabilize the condensate. Mg^2+^ therefore acts as a key regulator of RNA PS and can drive droplet formation of ssRNA at relatively low concentrations.^51,57^ The LCST behavior originates from the strengthening of Mg^2+^–phosphate interactions at elevated temperatures due to the entropy gain resulted from the releasing of hydration water. ^67^ Consistent with this mechanism, a set of control simulations, in which the Mg^2+^–phosphate potential is held fixed at 20 °C, leads to the dissolution of preformed condensates at high temperatures (Fig. S12), highlighting the importance of ion–RNA interactions in stabilizing the condensate.

**Figure 1.**
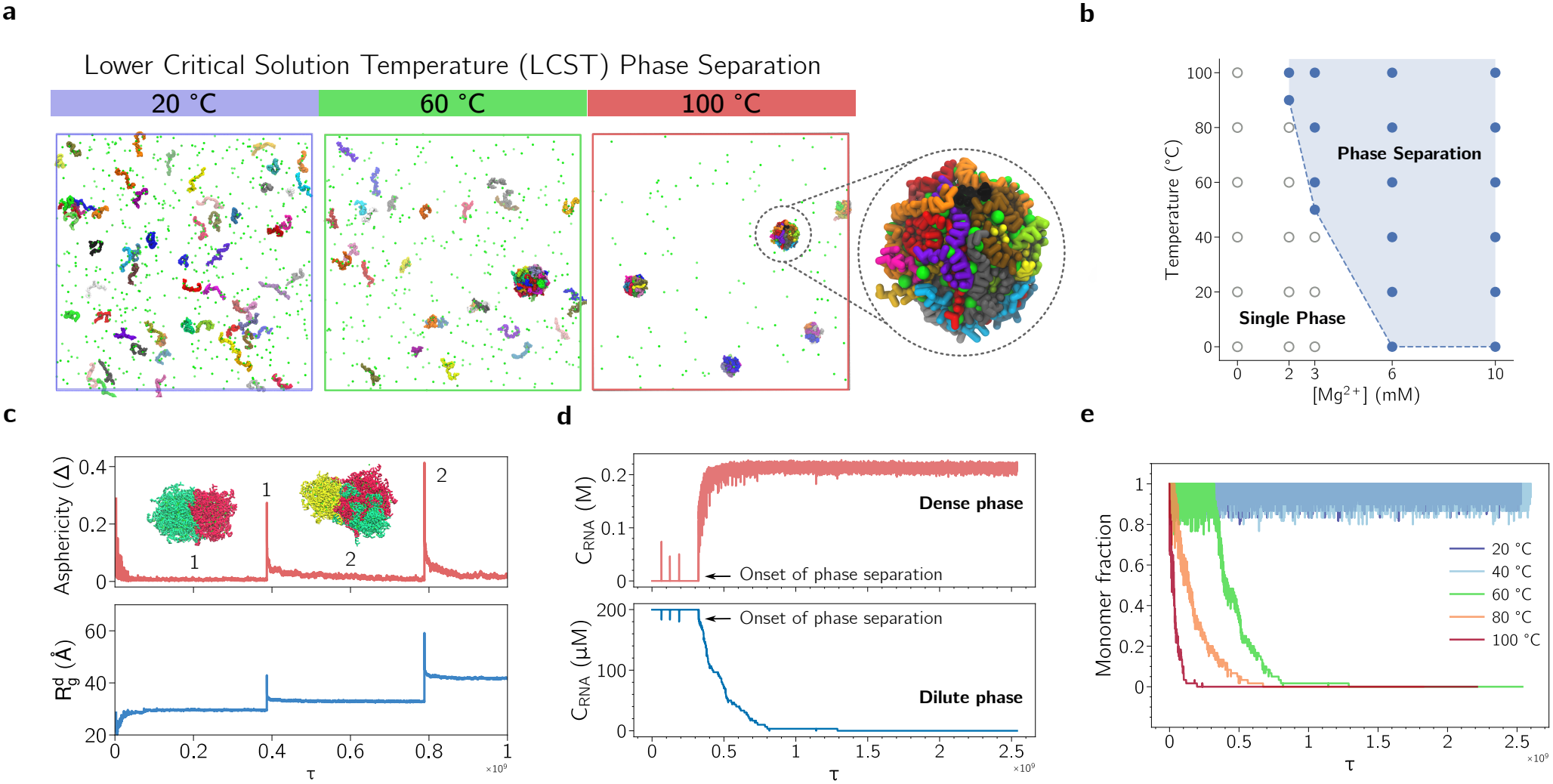
Ion- and temperature-mediated phase separation (PS) of RNA. **(a)** Temperature-dependent condensation of 200 *µ*M rA_30_ in 50 mM Na^+^ and 3 mM Mg^2+^ at 20, 60, and 100 °C, illustrating lower critical solution temperature (LCST) behavior. At elevated temperatures, RNAs transition from a dispersed dilute phase to a condensed phase. Chains are distinguished by different colors. **(b)** Putative phase diagram as a function of Mg^2+^ concentration and temperature. Filled circle indicate conditions where PS was observed from initially dispersed RNA chains, whereas open circles indicate conditions where no condensate was observed within the simulation timescale. **(c)** Time evolution of condensate asphericity (∆, top) and radius of gyration (*R*_*g*_, bottom), showing spontaneous droplet coalescence and the emergence of spherical condensates. Insets show representative snapshots of two coalescence events. **(d)** Time evolution of RNA concentration in the two phases at 3 mM Mg^2+^, demonstrating rapid enrichment of RNA in the dense phase (top) and depletion from the dilute phase (bottom). **(e)** Monomer fraction at different temperatures, showing accelerated PS at higher temperatures, consistent with LCST behavior.

The condensate remains near-spherical overall, indicated by an asphericity value close to zero (Fig. 1c). It is only disrupted by coalescence events, during which the condensate radius of gyration (R_g_) increases and its asphericity reaches ≈ 0.4, signifying a transiently oblate shape. In parallel, RNA concentration increases in the condensate while the dilute phase is depleted (Fig. 1d). Kinetically, PS occurs faster at higher temperatures or Mg^2+^ concentrations, where the monomer is depleted more rapidly (Figs. 1e, S3), consistent with the LCST behavior. In our simulations, the dense-phase RNA concentrations reaches 123 to 152 mM, corresponding to a nucleotide concentration of 3.7 to 4.6 M (Table S3, S4), which is comparable to measurements of ADP concentration in condensate (1.2 to 5 M). ^79^ Quantitative phase imaging (QPI)^80^ shows a 1000-fold increase of PGL-3 in condensate and 1000-fold increase of RNA concentration in FUS-RNA, which is in agreement with our simulations.

### Unstructured RNA collapses in condensate

Strikingly, we observe a pronounced collapse of RNA upon transitioning from the dilute phase to the condensate (Fig. 2a and S4). Specifically, the average R_*g*_ decreases from 23.00 ±0.03 to 17.50 ± 0.01 Å (Fig. 2a), and the average end-to-end distance, R_*ee*_, decreases from 60.0 ± 0.3 to 40 ± 1 Å (Fig. S4). Such a collapse is also confirmed by the Flory scaling exponent, *ν*, which quantifies the overall conformational state of a polymer.^81,82^ Typical values of *ν* correspond to distinct polymer conformations: *ν* = 1 for a rigid rod, *ν* ≈ 0.6 for a self-avoiding random walk (SAW), *ν* = 0.5 for a Gaussian random coil, and *ν* = 0.33 for a globule. To extract *ν*, we calculated the average pairwise distance, R(s), as a function of nucleotide separation *s* = |*i* − *j*|, where i and j denote nucleotide indices along the RNA and fit the tail of R(s) ∝ s^*ν*^. This analysis shows that *ν* decreases from ≈ 0.6 in the dilute phase to ≈ 0.5 in the condensate (Fig. 2b), indicating a transition from SAW-like conformations to Gaussian random-coil states upon RNA recruitment into condensate. Similar calculations using the form factor F(q)^83,84^ show the same behavior (Fig. S5). In addition, we find that RNA populates more spherical conformations in the condensate, as seen by the decrease in asphericity (Fig. S4).

**Figure 2.**
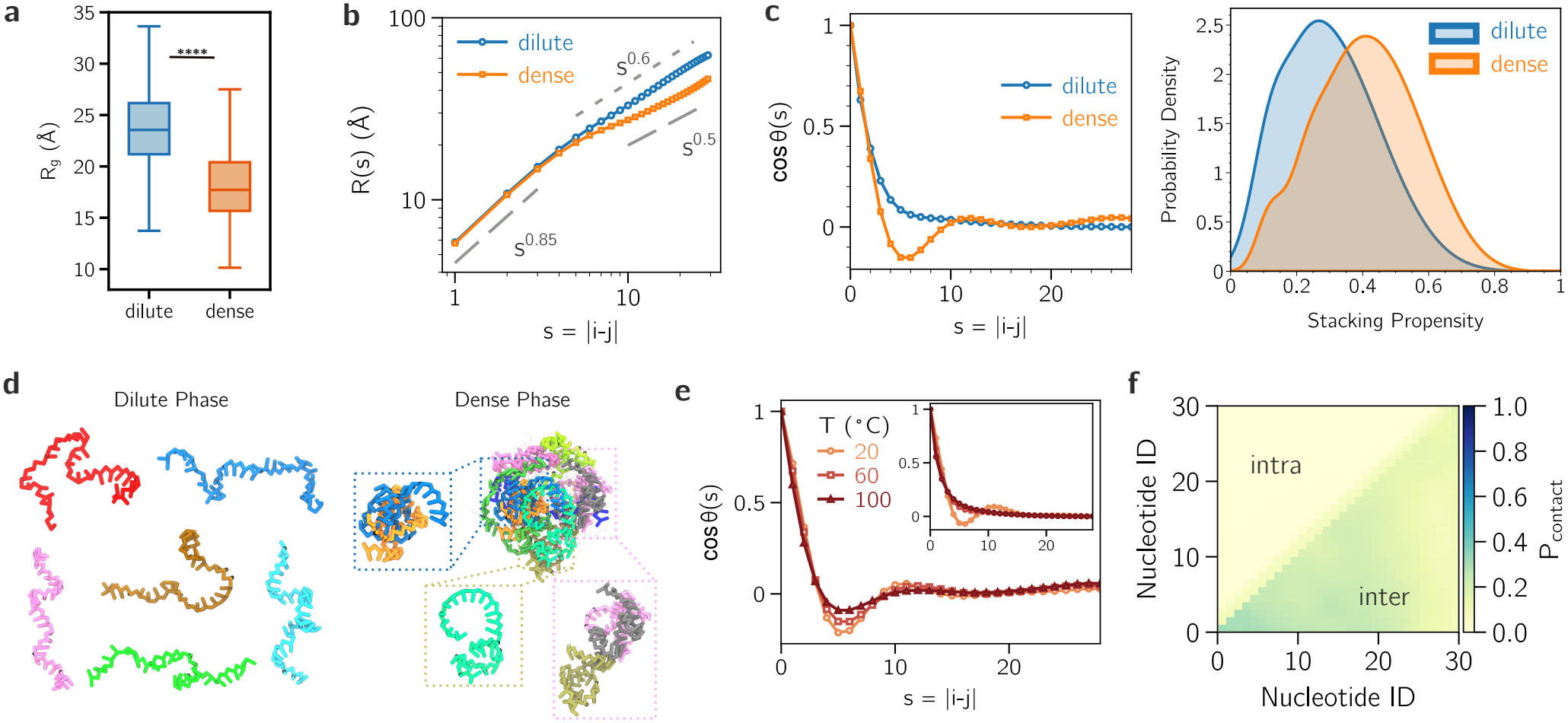
RNA structure in the two phases. The condition is for 200 *µ*M RNA at 300 mM Na^+^ and 3 mM Mg^2+^, unless specified otherwise. **(a)** Radius of gyration (R_*g*_) for rA_30_ in two phases at 20 °C. Asterisks indicate statistical significance (**** : p < 0.0001). Data were obtained from three independent simulation runs. Statistical significance was assessed using Welch’s t-test. For statistical analysis, each trajectory was divided into non-overlapping blocks of 5,000 frames, and the mean within each block was used as an independent observation. Uncertainties reported in the main text represent the standard deviation of the block-averaged values. **(b)** Mean internal distance, R(s), as a function of sequence separation *s* = |*i* − *j*| at 20 °C. Dashed and dotted lines denote SAW (*s*^0.6^) and Gaussian (*s*^0.5^) scaling, respectively. *s*^0.85^ corresponds to rod-like scaling due to the local stiffness. **(c)** Orientational correlation function (OCF), cos *θ*(*s*) vs. *s* (left), and stacking propensity (right) in the two phases at 60 °C. **(d)** Representative RNA conformations in the two phases. **(e)** OCF at different temperatures in the condensate. Inset: OCF in the dilute phase. **(f)** Intra and inter-chain contact map in condensate at 20 °C.

Our observation of unstructured RNA collapse in condensates is rare, as most previous studies have instead reported chain expansion upon condensate formation, including intrinsically disor-dered proteins (IDPs)^85–89^ and CAG-repeat RNAs.^18,41^ A recent simulation from Li and Chen predicted that rA_30_ would expand in condensate,^66^ contradicting our results. These differences reflect distinct modeling approaches to describing ion-mediated RNA interactions. However, from the polymer physics perspective, chains inside polymer melt should exhibit the Gaussian chain scaling (*ν* = 0.5) due to the similarity between intra- and interchain interactions. ^90,91^ Since rA_30_ is a polyelectrolyte which expands significantly in the dilute phase (*ν* ≈ 0.6), upon transitioning to the condensate, which is a rough approximation of a polymer melt, it should collapse to reduce *ν*. Indeed, recent high-speed atomic force microscopy experiments^92^ have revealed the collapse of poly-rA during condensate formation in the presence of Mg^2+^, supporting our findings. However, a recent SAXS experiment detected an increase of R_*g*_ (albeit with a reduction of *ν*) of those RNAs upon interacting with polypeptides in the dilute phase,^55^ suggesting proteins likely affect the RNA condensation process. Future experimental studies are therefore needed to resolve RNA conformations inside condensates.

Interestingly, the size and shape of RNAs in condensate show slight variations depending on the temperature and ionic conditions (Figs. S6, S7, S8). In particular, RNA becomes slightly more expanded with decreasing monovalent ion concentrations, presumably due to weaker electrostatic screening (Fig. S7). Upon heating, while R_*g*_ of the isolated chain shows a non-monotonic collapse,^67^ those inside condensates slightly expand (Fig. S6), suggesting a distinct environment in condensates. Together, these results indicate that although the overall conformational transition in the condensate is robust, the precise RNA structures might depend on various external factors. **ssRNA gains helicity in condensate**. Previous simulations^18,66,69^ and experiments^55,76,93,94^ have shown that the dense phase can significantly influence the nucleic acid structures. To assess structural changes between the two phases, we calculate the bond orientational correlation function (OCF), 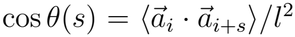(where 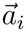 is the i^th^ bond vector and l is the bond length), which is often used to quantify the degree of chain helicity. ^55,95^ The correlation decays exponentially in the dilute phase, indicating that RNA lacks transient secondary structures (Figs. 2c and d). In contrast, in the condensate, we observe the emergence of clear oscillations in the OCF, which is a signature of partial helical ordering. Furthermore, the stacking propensity between consecutive bases increases from ≈ 0.25 in the dilute phase to ≈ 0.50 in the condensate (Fig. 2c). Together, these results indicate that rA_30_ gains helicity in the condensate through enhanced base-stacking interactions. Such a gain of partial structures has been documented for rU_30_ when interacting with polypeptides, while rA_30_ has been reported to lose its transient structures.^55^ At high temperatures, we observe that RNAs in the condensate largely retain their transient helical character while those in the dilute phase do not (Fig. 2e), suggesting that the crowded condensate environment strongly stabilizes stacked RNA conformations. Nevertheless, a slight decrease in the amplitude of the oscillations is observed with increasing temperature in the condensate (Figs. 2e and S9), presumably due to the weakening of stacking at elevated temperatures.

Despite its slight collapse in the condensate, we find that RNA intrachain contacts are few and distributed primarily along the diagonal of the contact map (Fig. 2f), which suggests that those contacts are largely local, with no pronounced long-range intrachain interactions. In contrast, interchain contacts are ≈ 4 times more frequent, suggesting that PS is primarily driven by intermolecular interactions. Interestingly, we find that the number of interchain contacts per chain, 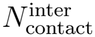, increases with temperature within the condensate (Fig. S10), further supporting the temperature-driven assembly process.

### LCST occurs due to stronger Mg^**2+**^accumulation and multidentate phosphate coordination

For monomers, our previous work^67^ showed that Mg^2+^ ions stabilize helical conformations by enhancing base-stacking interactions. Therefore, we hypothesize that the increased RNA helicity is due to the accumulation of Mg^2+^ in the condensate, which promotes base stacking and thus stabilizes RNA structures. To verify this hypothesis, we quantify Mg^2+^ local concentrations in condensate across a range of temperatures and ion concentrations. For this comparison, we report simulations with 300 mM Na^+^ in the background, as these systems undergo PS at all temperatures (20, 60, and 100 °C). Similar behaviors were observed at other monovalent ion concentrations.

Radial concentration profiles of Mg^2+^ from the condensate center-of-mass (COM) reveal strong enrichment of ions in the condensate relative to the dilute phase (Fig. 3a). These concentrations were calculated from the time-averaged number of Mg^2+^ ions in concentric spherical shells, nor-malized by their corresponding shell volumes (see SI Methods). Remarkably, those profiles exhibit peaks and valleys, indicating the existence of regions with enhanced or deplete ion accumulation. These nonuniform distribution are consistent with spatial heterogeneity of RNA packing and phos-phate density within the condensate, which can give rise to local difference in Mg^2+^ coordination and accumulation (Figs. 4h, i). As the temperature increases, Mg^2+^ becomes progressively depleted from the dilute phase and enriched in the condensate, suggesting enhanced uptake of Mg^2+^ by the RNA-rich phase (Fig. 3b). The Mg^2+^ concentrations in the two phases differ by more than three orders of magnitude, reaching 2 to 3 M in condensate, which is in qualitative agreement with experimental observations of complex coacervates formed by polyamines and nucleotides.^59^

**Figure 3.**
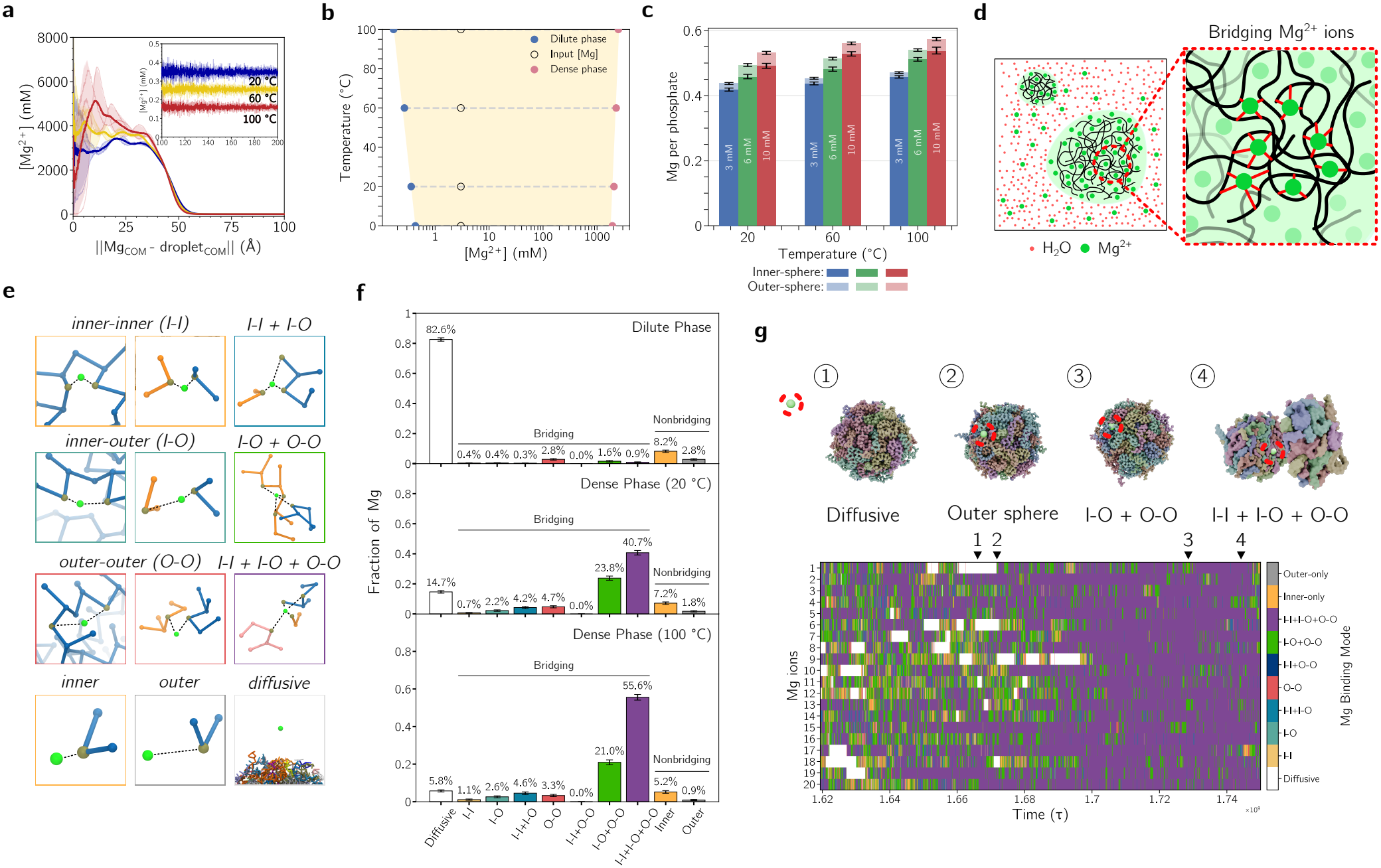
Temperature- and concentration-dependent Mg^**2+**^ **binding in rA**_**30**_ condensate. **(a)** Average Mg^2+^ radial concentration profiles from the condensate center of mass; inset, concentration in the dilute phase decreases with temperature. **(b)** Partitioning of Mg^2+^ between the two phases. **(c)** Number of Mg^2+^ per phosphate at various conditions partitioned into inner- and outer-sphere coordination. **(d)** Schematic of Mg^2+^ bridging interactions within the condensate. **(e)** Classification of Mg^2+^ binding modes, including bridging (I–I, I–O, O–O) and non-bridging (inner, outer, diffusive) states. **(f)** Fraction of Mg^2+^ in each binding mode in dilute (top) and dense (bottom) phases, showing enrichment of bridging states in the condensate. Error bars represent the standard deviation across trajectory blocks. **(g)** Representative trajectory of Mg^2+^ transitioning from diffusive to bound and bridging states. Snapshots illustrate coordination changes. Heat map shows binding-mode time series, where each row correspond to a distinct ion, with colors indicating specific binding modes.

**Figure 4.**
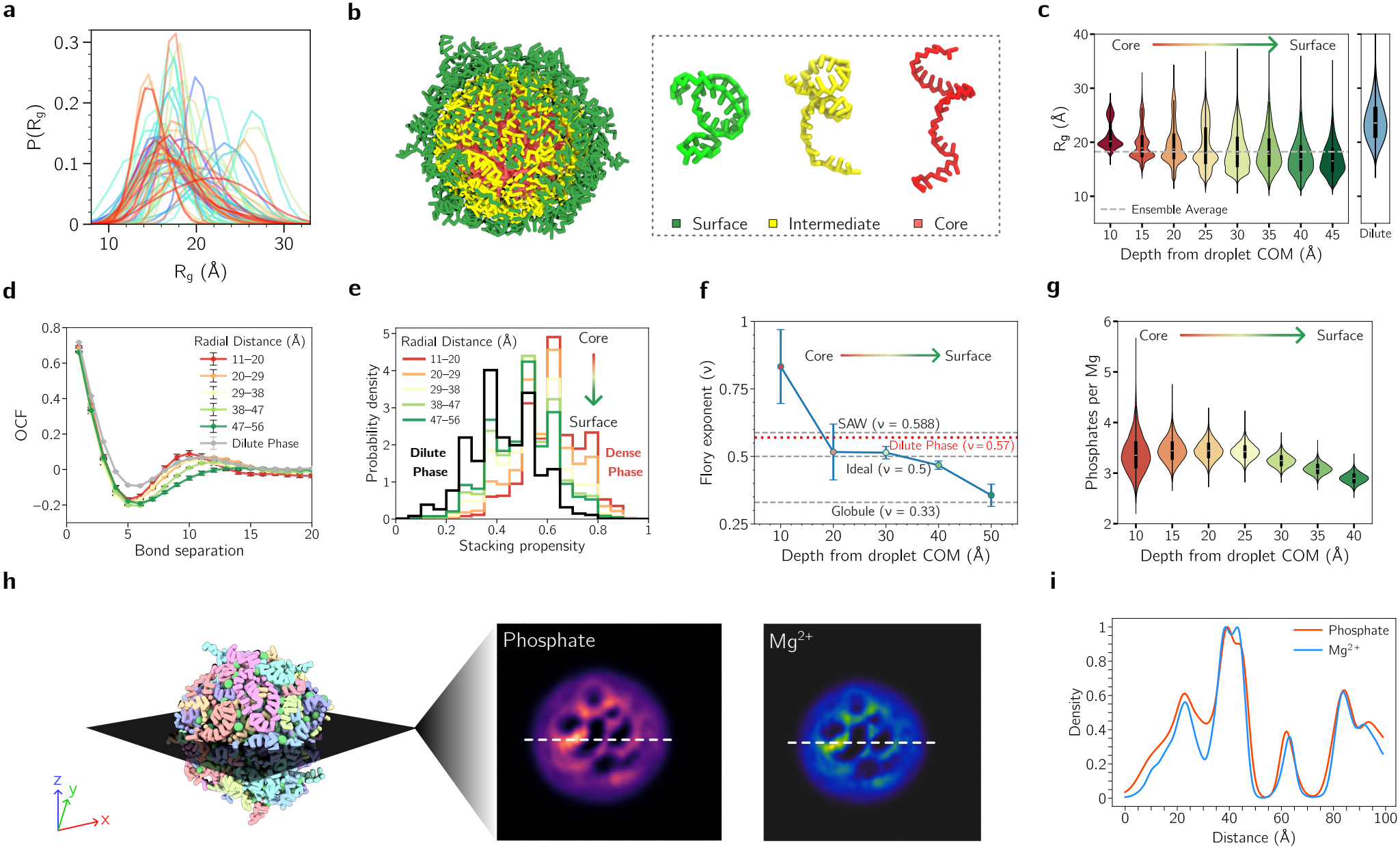
Heterogeneous network in RNA condensates mediated by Mg^**2+**^ ions. Unless otherwise stated, results are shown for 200 *µ*M rA_30_ at 300 mM Na^+^, 3 mM Mg^2+^ and 20 °C. **(a)** R_g_ for individual RNAs in condensate. **(b)** A snapshot of condensate with chains colored by radial position relative to the center of mass: core (red), intermediate (yellow), and surface (green). Also shown are representative conformations on the right. **(c)** R_g_ as a function of radial distance, showing expanded chains in the core and more compact chains near the interface. The dilute phase R_g_ is shown on the right for comparison. **(d)** Orientational correlation function for different layers of the droplet, indicating enhanced ordering in the condensate interior. **(e)** Base stacking propensity from condensate core to surface in 6 mM Mg^2+^ at 60 °C, showing stronger stacking in the core and reduced stacking near the interface. **(f)** Flory exponent decreases from self-avoiding walk (SAW)-like behavior in the condensate core (*ν* ≈ 0.6), through ideal-chain scaling (*ν* ≈ 0.5), to globule-like conformations (*ν* ≈ 0.3) near the condensate surface, indicating progressive chain collapse from the interior to the interface. The solid line represents the means across three independent simulation replicas, and error bars indicate 1 ±SEM. Reference lines indicated idealized polymer scaling regimes and the independently determined value for dilute phase. **(g)** Number of coordinated phosphates per Mg^2+^ as a function of radial distance, indicating increased bridging in the condensate interior. **(h)** Cross-sectional density maps of phosphate groups and Mg^2+^, showing co-localization and spatial heterogeneity (see also Supplementary Movie S1). **(i)** Radial density profiles of phosphates and Mg^2+^, demonstrating that they are co-localized.

To zoom in on how Mg^2+^ interacts with RNAs in condensate, we quantify Mg^2+^ coordination with phosphate groups by partitioning Mg^2+^ binding into inner- and outer-sphere modes, as done previously^70,96^ (Fig. 3c). Intriguingly, we found that the majority of Mg^2+^ in the condensate interacts using inner-sphere coordination, while only a small fraction binds through water-mediated interactions. This is reminiscent of a scenario where it is more favorable for Mg^2+^ to shed its hydration shell to bind directly to RNA due to the high electrostatic potential inside the RNA core.^96,97^ Increasing temperature mostly leads to an increase of inner-sphere coordination, while leaving outer-sphere population almost unchanged.

Due to the densely packed condensate, most ions bind to multiple phosphate groups at the same time (so-called bridging ions, Fig. 3d). We further partition them into inter-or intra-bridging depending on if the phosphate groups belonging to different or same RNA chains, respectively. In this scheme, Mg^2+^ may coordinate two phosphates through combinations of inner- and outer-sphere interactions, denoted as inner-inner (I-I), inner-outer (I-O), and outer-outer (O-O) states (Fig. 3e). A single Mg^2+^ ion can also form multiple bridges, such as I-I + I-O which indicates that it forms at least one inner-inner bridge and one inner-outer bridge. We find that the total population of bridging Mg^2+^ increases with both temperature and Mg^2+^ concentration (Figs. S11), supporting the ion-driven LCST behavior. In the dilute phase, Mg^2+^ ions are predominantly diffusive, accounting for 83% of the population; therefore, bridging is rare (only 6%) (Fig. 3f). In contrast, those in the condensate are mostly involved in bridging (76%), while the fraction of diffusive ions decreases sharply to 15%. Strikingly, the highly coordinated I-I + I-O + O-O state is the most dominant binding mode, contributing 41% of the total Mg^2+^ population. Increasing temperature further stabilizes most bridging modes while depleting the non-bridging and diffuse populations, suggesting that inter-chain bridging is the driving force for the LCST behavior of RNA, gluing different chains together by intermolecular crosslinks to stabilize the condensate. To further access the role of bridging ions, we performed control simulations in which Mg^2+^–phosphate interactions are held fixed at all temperatures. At 100 °C, a preformed condensate rapidly dissolves in this simulation, accompanied by a substantial reduction in the fraction of bridging Mg^2+^ ions from 40.1% to 3.4% (Fig. S12). This result demonstrates that the strengthening of Mg^2+^–phosphate interactions at high temperatures is crucial for maintaining the condensate by stabilizing multidentate coordination and interchain bridging networks.

Despite the high degree of ion coordination observed in condensates, we found that Mg^2+^ continuously exchanges between those binding states (Fig. 3g), similarly to the isolated monomers. ^98^ An excerpt shows the binding state of representative Mg^2+^ ions where each color denotes a distinct binding mode (Fig. 3g). This map reveals frequent transitions between different binding modes, indicating that Mg^2+^ coordination within the condensate is highly dynamic rather than locked into a single configuration.

### Network of heterogeneous RNA structures in condensate

Due to the large variation of Mg^2+^ concentration in the condensate, it is reasonable to suspect that the conformations of individual RNA chains might not be uniform. Instead, each chain could probably sample a distinct set of structures depending on its local ion concentration. Consistent with this idea, R_*g*_ values of individual chains span a broad distribution (Fig. 4a). The condensate is therefore stabilized by a heterogeneous network of interchain contacts:^41^ the interchain contact probability matrix reveals extensive, non-uniform patterns (Fig. S13) consistent with transient bridging and sequence-dependent local structures in condensates.

To dissect how local RNA structures vary within the condensate, we examine them as a function of radial positions relative to the condensate COM. Each RNA chain was assigned to a radial bin based on the position of its COM relative to the condensate COM. Therefore, these profiles describe the conformational properties of RNA chains centered at different radial positions, rather than those of individual nucleotides occupying that radial shell. ^86,87^ At first glance, chains in the core adopt more extended conformations, while those near the interface appear more compact (Fig. 4b). This is quantified by R_g_ as a function of radial distance, which decreases from the core to the interface (Figs. 4c, S22, S23). The Flory scaling further supports a pronounced conformational gradient across the condensate, where chains in the core exhibit self-avoiding walk (SAW)-like behavior (*ν* ≈ 0.6), transitioning toward ideal chain (*ν* ≈ 0.5) at intermediate depths, and finally becoming globule-like near the surface (*ν* ≈ 0.33) (Figs. 4f, S21). Furthermore, the OCFs show enhanced oscillations for chains in the core, but progressively weaken towards the interface, suggesting more stable RNA helical structures near the core (Fig. 4d). As a result, base stacking is also predominant in the core and decreases toward the interface (Fig. 4e). Together, these radial gradients in chain size and secondary structure indicate that the condensate interior stabilizes more expanded ssRNA conformations alongside enhanced base-base stacking to form transient RNA helical structures. This is in line with our previous simulations of CAG-repeat RNA condensates showing that recruitment into the dense phase drives a hairpin-to-extended transition that builds a system-spanning, intermolecular RNA network. ^18,41,99^ It is also supported by recent experiments and simulations proposing that condensates can undergo percolation transitions enabled by RNA-specific base stacking/base pairing and ionic interactions that potentially lead to dynamical arrest.^51,54,57,100^ Similar behavior has also been found for proteins in simulations with a much larger system size,^85,87,101^ suggesting a general mechanism of heterogeneity in condensates.

### Mg^2+^-mediated bridging reinforces the condensate interior

Due to such crowded environment, the number of phosphates coordinated per Mg^2+^ ion in the core is highest and decreases gradually to the interface (Fig. 4g). This suggests that the condensate core engages in more extensive multi-phosphate coordination and thus multivalent bridging. Mg^2+^ therefore acts not only as screening ions but as crosslinkers that couple multiple phosphate groups and thus contribute to a correlated ionic network. This is consistent with recent simulations showing that RNA PS emerges from the interplay of Mg^2+^-phosphate interactions, stacking, and hydrogen bonding,^66,68^ as well as experiments showing that RNA condensation is driven by heterotypic RNA-Mg^2+^ interactions.^51,57,102^ Consistent with this picture, the cross-sectional density maps from our simulations reveal strong spatial co-localization of Mg^2+^ and phosphate groups, with both exhibiting heterogeneous organization within condensate (Fig. 4h, Supplementary Movie S1). The density profiles show that the two closely follow each other (Fig. 4i), indicating that ion enrichment is tightly coupled to RNA density. Comparing the first and second halves of the simulated trajectories shows consistent radial concentration profiles (Figs. S24 to S27), supporting the stability of those spatial distributions over time.

### Sequence dependence of ion accumulation in condensate

RNA sequence affects condensate morphology, dynamics, and material properties, in part through differences in base stacking, base pairing, and ion-mediated interactions. ^16,49,103–105^ Purine-rich RNAs generally have a greater propensity to form stacked or transiently ordered conformations, whereas pyrimidine-rich RNAs are less locally structured.^55,67,95,106^ To examine how these sequence-dependent interactions influence condensate organization, we next compare rA_30_ and rU_30_ in both dilute and condensed phases.

In the dilute phase, rA_30_ and rU_30_ exhibit similar sizes.^67^ Inside condensates, however, rA_30_ adopts slightly more compact conformations than rU_30_, with an average decrease of 1.0 Å in chain size (Figs. 5b and S14). This is consistent with the greater propensity of rA_30_ to form transient helical and base-stacked structures, as reflected by the more pronounced oscillatory behavior of its OCF (Fig. 5b).^67,95^ The magnitude of such differences depends on both temperature and Mg^2+^ concentration (Fig. S15). At low temperatures and Mg^2+^ concentrations, rA_30_ is generally more compact than rU_30_. At other conditions, however, this sequence-dependent difference becomes less pronounced, leading to similar chain dimensions or, in some cases, slightly larger rA_30_ conformations relative to rU_30_, presumably due to the former’s larger base.

**Figure 5.**
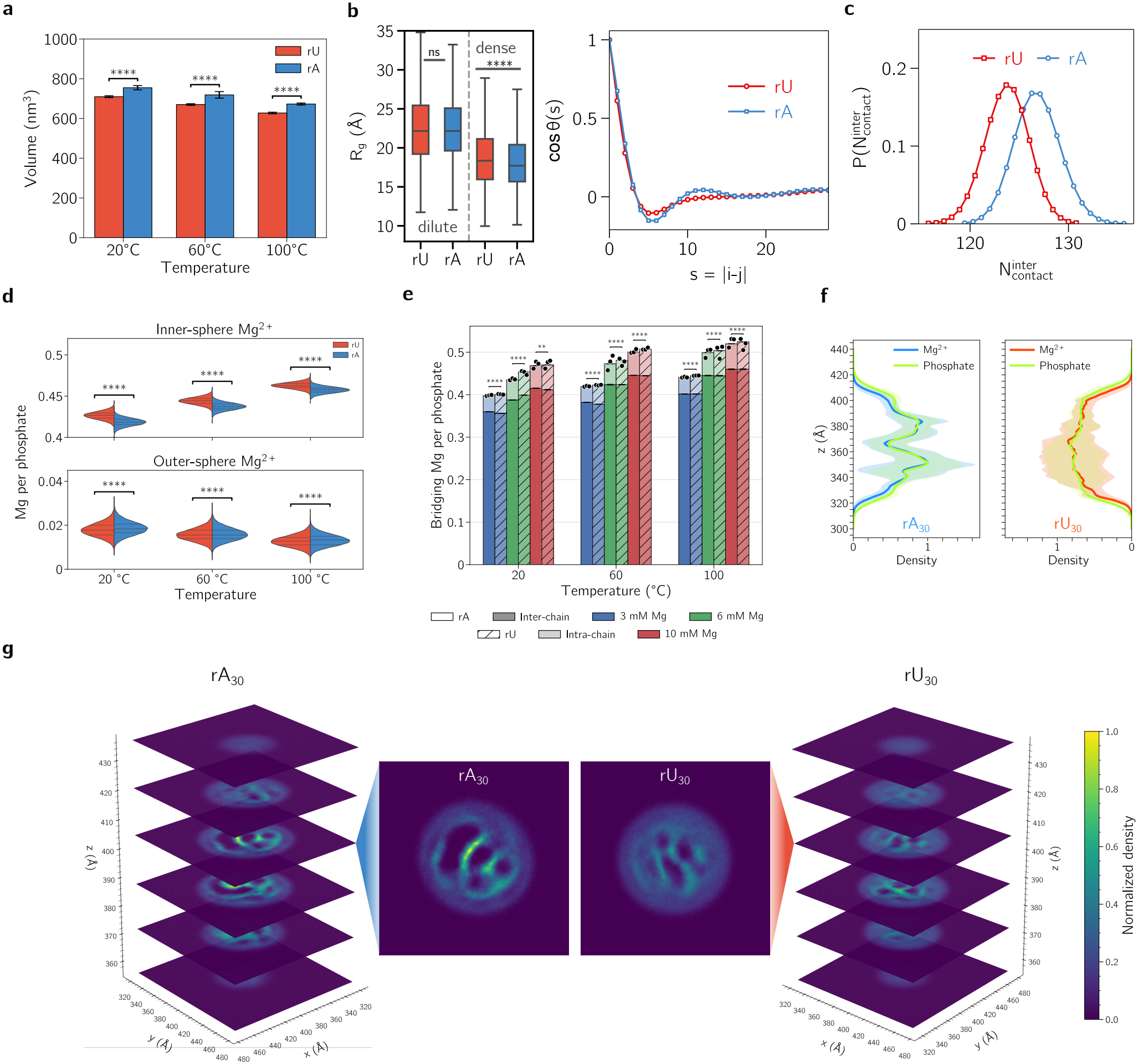
Effect of sequence on condensate properties. Unless otherwise noted, all results are at 300 mM Na^+^ and 3 mM Mg^2+^. **(a)** Volume of rA_30_ and rU_30_ condensates at 20, 60, 100 °C. **(b)** Left: *R*_*g*_ for individual chains in the two phases at 20 °C. Right: Bond-vector orientational correlation function, showing enhanced local structures for rA_30_ relative to rU_30_ in condensate. **(c)** Number of interchain contacts per RNA. **(d)** Distribution of inner- and outer-sphere Mg^2+^ coordination per phosphate for rA_30_ and rU_30_ condensates at 20, 60, and 100 °C. **(e)** Number of bridging Mg^2+^ per phosphate for inter- and intrachain coordination at varying conditions. **(f)** One-dimensional density profiles along the z-axis obtained from cylindrical slices with radius of 10 Å at 20 °C. Solid lines denote the average normalized density, and shaded regions indicate the standard deviation. **(g)** Stacked Mg^2+^ density slices along the z-axis, with representative slices at 100 °C (see also Supplementary Movie S2).

Despite being more compact at the individual chain level, surprisingly rA_30_ condensates are slightly larger across all temperatures (Fig. 5a). This indicates that the condensate volume is not determined solely by the size of individual RNA chains. Instead, dense-phase packing likely depends on how RNA chains, ions, and local structural motifs are collectively organized. One possibility is that rU_30_ aligns better to pack more efficiently in the condensate. However, we did not observe any differences in the orientational packing of RNA chains between the two using the nematic order parameter, which quantifies the degree of alignment in anisotropic systems. Both rA_30_ and rU_30_ exhibit similarly low nematic order values (Fig. S16), indicating weak alignment.

We then asked if sequence-dependent differences in local RNA contacts and Mg^2+^ organization contribute to the packing in condensate. Short-range RNA-RNA radial distribution functions are broadly similar for rA_30_ and rU_30_, with only modest sequence-dependent differences in peak intensity (Fig. S17). Interestingly, we find that rA_30_ condensates form more P-P contacts than rU_30_ (Figs. 5c, S18) despite being slightly larger, which indicates that the larger rA_30_ condensate does not reflect weaker local RNA association. Instead, in rA_30_ condensates, RNA chains are concentrated in more localized high-density regions, suggesting a more spatially heterogeneous distribution within condensate. In contrast, rU_30_ displays a more homogeneous distribution in condensate (Figs. 5g, S28, S29, S30, Supplementary Movie S2). Together, these results show that rA_30_ and rU_30_ adopt distinct organizations in condensate: rA_30_ condensates are characterized by local RNA-rich clustering and base-stacking associated with chain collapse, whereas those formed by rU_30_ exhibit more uniform organization with efficient packing.

Because Mg^2+^ plays a central role in screening RNA charge and bridging phosphate groups, we next examine whether sequence-dependent Mg^2+^ coordination contributes to those differences in condensate organization. Remarkably, rU_30_ exhibits a stronger preference for inner-sphere Mg^2+^ coordination in condensate, whereas rA_30_ contains a slightly larger fraction of outer-sphere Mg^2+^ (Fig. 5d). Higher temperatures shift the binding from outer-to inner-sphere, supporting the ion-induced LCST behavior. Due to the higher preference for inner-shell ion binding and its smaller volume, rU_30_ condensate contains slightly more bridging Mg^2+^ across all conditions. These bridging ions may, therefore, facilitate more uniform electrostatic coupling between RNAs, reducing local heterogeneity and enabling denser packing in rU_30_ condensates. In support of this interpretation, cylindrical averaging of the density reveals that rA_30_ condensate exhibits pronounced spatial fluctuations, characterized by jagged density profiles across the cross section. In contrast, rU_30_ condensate displays a smoother density profile, consistent with a more homogeneous organization. The phosphate profile closely follows those of Mg^2+^, indicating closely coupled spatial distributions of RNA and Mg^2+^ within the condensate (Fig. 5f). Future experiments, such as SAXS^55^ and FRET,^107^ could provide insights into RNA conformational changes within condensates and help assess our predictions regarding RNA sequence dependent properties upon condensate formation.

## Discussions and Conclusion

Our simulations reveal that unstructured RNAs within condensates surprisingly adopt transient helical conformations that are much less pronounced in the dilute phase. This is in contrast with previous studies^17,18,41,66,108^ suggesting that other sequences such as CAG or CUG repeats, which form hairpin-like conformations in the dilute phase, can lose their secondary structures in condensates. If verified, structural transitions within condensates thus depend significantly on the intrinsic structure of RNA. Our finding is supported by a recent SAXS experiment where rU_30_ has been shown to populate transient helical structures in RNA-protein complexes right before condensation.^55^ It suggests that condensates may promote disorder-to-order transitions for unstructured RNAs while inducing the opposite behavior for folded RNAs, thereby acting as structure-promoting or structure-disrupting environments depending on the sequence context. Such structural remodeling may influence protein and ion/metabolite recruitment within condensates. For example, RNA-binding proteins (RBPs) with structured RNA-binding domains are often associated with recognition of helical RNAs, whereas intrinsically disordered RBPs may preferentially interact with less structured RNA regions. ^109,110^ Therefore, the gain or loss of RNA structure within condensates may serve as a key determinant of protein selectivity, ultimately shaping the composition and function of cellular condensates. Similarly, recent computational studies^88,111^ have shown that condensates can also remodel protein conformational ensembles depending on their environment and the interactions between client and scaffold proteins. Taken together, our findings further highlight the role of the condensate environment in dictating changes in molecular conformations.

Interestingly, we find that RNA structure in condensates is not uniform, similar to the CAG-repeat.^18,41^ Instead, RNA conformations are spatially heterogeneous, with molecules in the core adopting more extended and helical structures, while those near the surface are comparatively less helical and more compact. In addition, the ion distribution within condensates is also inherently heterogeneous. Such structural heterogeneity and microenvironmental diversity may have important implications for the spatial organization of biomolecules within condensates and the emergence of nanoscale domains^32,38,112^ and multiphase architectures^2,37,41^ with distinct physico-chemical conditions and material properties. Indeed, many cellular condensates, such as nucleoli, stress granules, and paraspeckles, exhibit multilayered organization.^5,29,30^ For example, the nucleolus is partitioned into multiple layers, each associated with distinct stages of ribosome biogenesis, including rDNA transcription, pre-rRNA processing, and ribosome assembly.^5^ These compart-mentalized functions require selective enrichment of specific proteins within different subdomains. In this context, spatial variation in RNA structure could contribute to differential protein localization within condensates. For instance, proteins such as NONO and FUS have been reported to preferentially associate with structured regions of NEAT1 RNA and localize to the core of paraspeckles, whereas proteins like TDP-43 tend to interact with less structured regions and are enriched near the periphery.^29^ Such observations suggest that spatial structural heterogeneity of RNA within condensates may contribute to the organization of membraneless compartments into distinct subphases by guiding selective protein recruitment. This, in turn, can support functional specialization within condensates and give rise to complex internal organization, with potential implications for early prebiotic compartmentalization and the origin of life.^113^

In addition to RNA structural changes, we find that condensates accumulate thousands-fold higher concentrations of Mg^2+^ compared to the bulk solution. Mg^2+^ ions are known to play important roles in stabilizing the secondary and tertiary structures of biomolecules through ion binding and bridging interactions. ^70,71,84,114–118^ Therefore, such elevated ion concentrations within condensates may create a distinct environment that favors the folding of biomolecules and helps stabilize their functionally active conformations. Consistent with this view, previous studies have suggested that condensates can influence the stability of RNA and proteins.^44,69,76–78,119^ This raises the possibility that condensates may act as chaperone-like environments that facilitate RNA folding and subsequent catalytic activity.^22,75^ In addition, the accumulation of ions in condensates establishes a gradient between the two phases, generating an electrochemical potential across the condensate boundary.^33,34,42–44,120,121^ Such a potential may influence the partitioning of charged species, thereby modulating reaction rates within condensates. ^36,42,75,122^ It can also facilitate directed transport of charged molecules within the cellular environment, contributing to molecular flux and spatial organization.^123–125^

In addition to electrostatic-mediated interactions, recent computational studies^126,127^ have shown that the competition between electrostatic interactions and sequence-dependent molecular stickiness influences the selective partitioning of ssRNA and dsRNA into protein–RNA condensates. Similarly, the selective partitioning of intrinsically disordered regions (IDRs) into protein condensates depends on the balance between client–scaffold interactions and scaffold self-interactions. Thus, selective molecular partitioning in RNA condensates is likely governed by multiple factors, including condensate composition, client and scaffold sequences, and the local microenvironment. Understanding how these factors collectively regulate molecular partitioning remains an important direction for future research.

In conclusion, our work demonstrates that RNA-driven condensates exhibit remarkable structural and ionic heterogeneity, even when formed via homotypic RNA–RNA interactions mediated by multivalent ions. Temperature and ion concentration jointly modulate RNA structures, ion partitioning, and condensate properties, giving rise to sequence-dependent architectures and spatially distinct microenvironments. These features mirror the multilayered organization observed in cellular condensates and provide insights into selective biomolecule recruitment, enzymatic regulation, and condensate aging. Importantly, our findings highlight design principles for synthetic biomaterials: by tuning RNA sequence, ion identity, ion concentration, and temperature, it is possible to engineer condensates with programmable internal architecture and chemical microenvironments. Such rationally designed condensates could serve as functional compartments for controlled molecular partitioning, catalytic activity, and drug delivery, offering a framework to translate the principles of RNA phase separation into next-generation biomaterials.

## Methods

### RNA energy function

To simulate temperature-dependent RNA phase separation, we utilized the three-interaction-site RNA model with temperature-dependent Mg^2+^−P interaction potential, which predicts the temperature induced RNA compaction.^67^ All details can be found in the original work.^67,70^ Briefly, each nucleotide is represented by three beads, corresponding to the phosphate, ribose, and nucleobase. The beads are positioned at the COM of the chemical groups. The energy function for an isolated RNA chain has the following form:

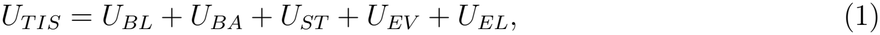

which describes the interactions between RNA beads as a sum of contributions from the bond length (U_*BL*_), bond angle (U_*BA*_), single-stranded stacking (U_*ST*_), excluded volume (U_*EV*_), and electrostatic (U_*EL*_) interactions. We used the harmonic approximation for U_*BL*_ and U_*BA*_. The excluded volume interactions U_*EV*_ is given by the Weeks-Chandler-Andersen potential (WCA):^128^

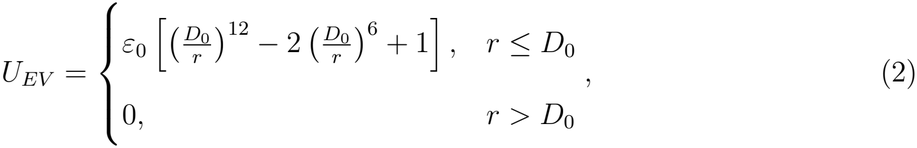

where ε_0_ = 1 kcal/mol and D_0_ = R_*i*_ + R_*j*_. The WCA term vanishes if the interacting sites are separated by a distance greater than D_0_. For interactions between two bases, we set D_0_ = 3.2 Å to favor stacking.

The stacking interaction is modeled using

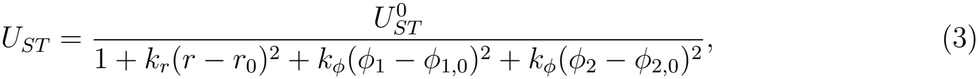

where k_*r*_ and k_*ϕ*_ are the spring constants, r is the distance between two consecutive bases, ϕ_1_(P_*i*_, S_*i*_, P_*i*+1_, S_*i*+1_) and ϕ_2_(P_*i*+2_, S_*i*+1_, P_*i*+1_, S_*i*_) are the dihedral angles among phosphate-sugar beads. These parameters are taken from an A-form RNA helix. 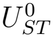 is optimized from the melting of different RNA dinucleotides. The details of the parameters are provided elsewhere. ^67,70^ Following our previous procedure,^67^ we employ the one-dimensional Reference Interaction Site Model (1D-RISM) to compute the Mg^2+^–phosphate potentials of mean force (PMF) at the temperatures of interest. This approach provides an efficient framework to incorporate solvent-mediated ion–phosphate interactions, including short-range ion correlations and temperature-dependent dielectric effects, without requiring explicit solvent simulations.

Within this framework, 1D-RISM yields the radial distribution function between Mg^2+^ ions and phosphate groups, g_Mg*−*P_(r), from which the PMF is obtained as:

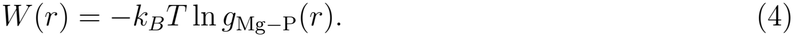

The electrostatic interaction term (U_*EL*_) consists of Debye-Hückel (DH) and a temperature-dependent potential of mean force (PMF). Electrostatic effects from monovalent ions are treated implicitly using DH potential while interactions between Mg^2+^ and phosphates are captured through the PMF using:

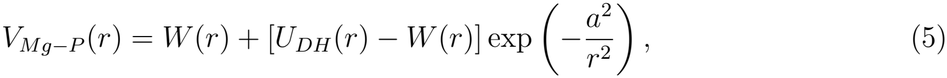

which ensures the correct long-range behavior by smoothly merging the short-range PMF, W (r), with the mean-field DH potential (U_*DH*_). The constant a = 5.0 Å preserves the Mg^2+^−P direct interaction energy. The resulting effective potential V_*Mg−P*_ (r) asymptotically reaches the DH approximation in long range, while displaying water-mediated short-ranged interactions according to RISM theory.

Electrostatic interactions between phosphate–phosphate and Mg^2+^–Mg^2+^ are treated using DH theory:

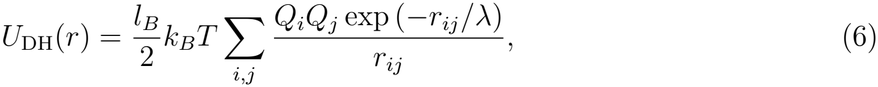

where λ = (8πl_*B*_ρ_1_)^*−*1*/*2^ is the Debye screening length determined by the number density of monovalent ions ρ_1_. The Bjerrum length,

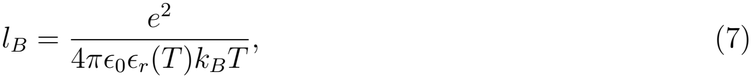

sets the strength of electrostatic interactions. Here, e is the elementary charge, ϵ_0_ is the vacuum permittivity, and ϵ_*r*_(t) is the temperature-dependent dielectric constant of water:

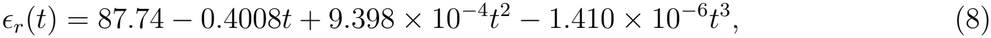

where t is in °C.

Finally, the effective phosphate charge Q_*P*_ (T, C_1_, C_2_) depends on both monovalent (C_1_) and di-valent (C_2_) ion concentrations via counterion condensation theory (see details in Refs. ^67,70^). This treatment renormalizes the bare phosphate charge to account for ion-mediated screening and condensation effects, providing an effective charge consistent with the ionic environment.

### Intermolecular Interactions

In our model, direct interactions between RNA chains consist of excluded volume repulsion described by the WCA potential and screened eletrostatic repulsion between phosphates described by Debye-Hückel theory. Base-stacking interactions are restricted to consecutive nucleotides within the same RNA chain, and no hydrogen bonding interactions are included. Therefore, the model does not contain direct attractive interactions between RNA chains. Instead, interchain association is mediated by explicit Mg^2+^ ions interacting with RNA phosphate groups through a temperatue-dependent potential derived from 1D-RISM calculations. The Mg^2+^–phosphate potential is parameterized separately at each temperature and becomes more attractive upon heating, favoring multidentate phopshate coordination and Mg^2+^-mediated interchain bridging. In addition, the electrostatic interactions depend on temperature through the dielectric constant, Bjerrum length, and effective phosphate charge. Thus, the observed LCST-type phase separation reflects the temperature dependence incorporated into the effective ion–RNA interaction model.

## Acknowledgments

We thank Phil Bevilacqua for critical reading of the manuscript. We gratefully acknowledge financial support from the University at Buffalo to H.T.N.. Computational resources were provided by the Center for Computational Research at the University at Buffalo.

## Data and Software Availability

The data underlying this study are available in the published article, the Supporting Information, and at https://zenodo.org/records/21839533. The simulation code is available at https://github.com/BioSimLabUB/biophysical-temperature-dependent-ssRNA.

