## Supplementary Information for "Molecular simulation of ion-mediated phase separation and structural organization in RNA condensates"

Lipika Baidya,[†,‡] Heyang Zhang,[†,‡] and Hung T. Nguyen[*,†,¶]

†*Department of Chemistry, State University of New York at Buffalo, Buffalo, New York 14260-3000, United States*

‡*These authors contributed equally to this work.*

¶*Institute for Artificial Intelligence & Data Science, State University of New York at Buffalo, Buffalo, NY, USA*

**Tables** **15**

**Figures** **18**

**4 Effect of monovalent ions on phase separation** **19**

**References** **46**

### 1 RNA Model

To simulate temperature-dependent RNA phase separation, we utilized the three-interaction-site RNA model with temperature-dependent $Mg^{2+}$−P interaction potential, which predicts the temperature induced RNA compaction.[1] All details can be found in the original work.[1,2] Briefly, each nucleotide is represented by three beads, corresponding to the phosphate, ribose, and nucleobase. The beads are positioned at the center of mass of the chemical groups. The energy function for an isolated RNA chain has the following form:

$$U_{TIS} = U_{BL} + U_{BA} + U_{ST} + U_{EV} + U_{EL}, \tag{1}$$

which describes the interactions between RNA beads as a sum of contributions from the bond length ($U_{BL}$), bond angle ($U_{BA}$), single-stranded stacking ($U_{ST}$), excluded volume ($U_{EV}$), and electrostatic ($U_{EL}$) interactions. We used the harmonic approximation for $U_{BL}$ and $U_{BA}$. The excluded volume interactions $U_{EV}$ is given by the Weeks-Chandler-Andersen potential (WCA):[3]

$$U_{EV} = \begin{cases} \varepsilon_0 \left[ \left(\frac{D_0}{r}\right)^{12} - 2\left(\frac{D_0}{r}\right)^{6} + 1 \right], & r \le D_0 \\ 0, & r > D_0 \end{cases}, \tag{2}$$

where $\varepsilon_0 = 1$ kcal/mol and $D_0 = R_i + R_j$. The WCA term vanishes if the interacting sites are separated by a distance greater than $D_0$. For interactions between two bases, we set $D_0 = 3.2$ Å to favor stacking.

### 2 Electrostatic interactions

#### $Mg^{2+}$–P PMFs

Following our previous procedure,[1] we employ the one-dimensional Reference Interaction Site Model (1D-RISM) to compute the $Mg^{2+}$–phosphate potentials of mean force (PMF) at the temperatures of interest. This approach provides an efficient framework to incorporate solvent-mediated ion–phosphate interactions, including short-range ion correlations and temperature-dependent dielectric effects, without requiring explicit solvent simulations.

Within this framework, 1D-RISM yields the radial distribution function between $Mg^{2+}$ ions and phosphate groups, $g_{\mathrm{Mg-P}}(r)$, from which the PMF is obtained as:

$$W(r) = -k_B T \ln g_{\mathrm{Mg-P}}(r). \tag{3}$$

#### Effective $Mg^{2+}$–P interaction potential

The electrostatic interaction term ($U_{EL}$) consists of Debye-Hückel (DH) and a temperature-dependent potential of mean force (PMF). Electrostatic effects from monovalent ions are treated implicitly using DH potential while interactions between $Mg^{2+}$ and phosphates are captured through the PMF using:

$$V_{Mg-P}(r) = W(r) + [U_{DH}(r) - W(r)] \exp\left(-\frac{a^2}{r^2}\right), \tag{4}$$

which ensures the correct long-range behavior by smoothly merging the short-range PMF, $W(r)$, with the mean-field DH potential ($U_{DH}$). The constant $a = 5.0$ Å preserves the $Mg^{2+}-P$ direct interaction energy. The resulting effective potential $V_{Mg-P}(r)$ asymptotically reaches the DH approximation in long range, while displaying water-mediated short-ranged interactions according to RISM theory.

##### Electrostatics for other interactions

Electrostatic interactions between phosphate–phosphate and $Mg^{2+}$–$Mg^{2+}$ are treated using DH theory:

$$U_{\mathrm{DH}}(r) = \frac{l_B}{2} k_B T \sum_{i,j} \frac{Q_i Q_j \exp\left(-r_{ij}/\lambda\right)}{r_{ij}}, \tag{5}$$

where $\lambda = (8\pi l_B \rho_1)^{-1/2}$ is the Debye screening length determined by the number density of monovalent ions $\rho_1$. The Bjerrum length,

$$l_B = \frac{e^2}{4\pi\epsilon_0\epsilon_r(T)k_BT}, \tag{6}$$

sets the strength of electrostatic interactions. Here, $e$ is the elementary charge, $\epsilon_0$ is the vacuum permittivity, and $\epsilon_r(t)$ is the temperature-dependent dielectric constant of water:

$$\epsilon_r(t) = 87.74 - 0.4008t + 9.398 \times 10^{-4}t^2 - 1.410 \times 10^{-6}t^3, \tag{7}$$

where $t$ is in °C.

Finally, the effective phosphate charge $Q_P(T, C_1, C_2)$ depends on both monovalent ($C_1$) and divalent ($C_2$) ion concentrations via counterion condensation theory (see details in[1,2]). This treatment renormalizes the bare phosphate charge to account for ion-mediated screening and condensation effects, providing an effective charge consistent with the ionic environment.

### 3 Simulation details and analyses

#### 3.1 Simulation details

Sixty fully extended RNA chains ($rA_{30}$ or $rU_{30}$) were placed evenly in a cubic simulation box with side length of 793 Å for RNA concentrations of 200 $\mu$M. The number of $Mg^{2+}$ ions were chosen to exceed at least half the number of phosphate groups (900-3000 ions depending on the ion concentrations), ensuring a sufficient population of ions in the dilute phase after phase separation occurs. All simulations were performed using OpenMM[4] (version 8.1.0) on NVIDIA A100 graphics processing units. The equations of motion were integrated in the NVT ensemble using the LangevinMiddleIntegrator[5] with a time step of 2 fs. To enhance the conformational sampling, low-friction Langevin dynamics were employed with a friction coefficient of 0.1 $ps^{-1}$. Periodic boundary conditions were applied in all three spatial dimensions to minimize finite-size effects. Simulation snapshots were recorded every 10,000 steps. All trajectories were examined for equilibration using the timeseries analysis implemented in PyMBAR[6] as done previously.[1] The equilibration time $t_0$ was determined by maximizing the number of effectively uncorrelated samples, thereby identifying the optimal production window. Data prior to $t_0$ were discarded from all analyses. Statistical convergence was further evaluated by verifying the stability of key observables (e.g. $R_g$, ion counts) over time, and by dividing each trajectory into two equal halves and confirming consistent ensemble averages between them.

To construct the phase diagrams, simulations were performed at $Mg^{2+}$ concentrations of 0, 2, 3, 6, 10 mM; $Na^+$ concentrations of 50 and 300 mM; and temperatures of 0, 20, 40, 80, and 100 °C. Each simulation was conducted for 2 to 10 $\mu$s to access the presence or absence of phase separation. Phase behavior was classified based on the formation and persistence of RNA-rich condensates coexisting with a dilute phase.

#### 3.2 Temperature-independent $Mg^{2+}$–phosphate interaction simulation

To access the role of the temperature-dependent $Mg^{2+}$–phosphate potential of mean force (PMF) in stabilizing RNA condensates, we performed an additional control simulation of $rA_{30}$ at 200 $\mu$M RNA, 50 mM $Na^+$, and 3 mM $Mg^{2+}$. The control simulation was initialized from the final snapshot of a trajectory containing a preformed condensate of 60 RNA chains obtained at 100 °C using the $Mg^{2+}$–phosphate PMF parameterized at 100 °C. The $Mg^{2+}$–phosphate interaction potential was then replaced with its 20 °C counterpart, while the simulation temperature was still held at 100 °C. The control simulation was performed for 1.2 $\mu$s with a time step of 2 fs. Under the modified potential, the condensate quickly dissolved, accompanies by substantial reduction in the fraction of bridging $Mg^{2+}$ ion from 49.1 % in the original system to 3.4 % in the control.

#### 3.3 Convergence analysis and uncertainty estimation

To evaluate the temporal stability of radial concentration and RNA conformation profiles, the analyzed production trajectories of each independent replica was divided into two non-overlapping halves. Radial phosphates and $Mg^{2+}$ concentration profiles were calculated separately for each half and averaged across three independent replicas. Pearson correlation coefficients between the first- and second-half concentration profiles were calculated independently for each replica to evaluate similarity between the two halves.

#### 3.4 Clustering

For each snapshot, RNA chains were grouped into clusters based on a distance-based connectivity criterion. Two RNA monomers were considered part of the same cluster if any pair of nucleotides, one from each monomer, were within 10 Å of each other, corresponding to the excluded-volume cutoff distance in our model. Cluster membership was defined recursively: a monomer was assigned to a cluster if it satisfied this distance criterion with any monomer already in that cluster.

##### 3.5 Radius of gyration and shape parameters

$R_g$ and shape descriptor were calculated from the gyration tensor defined as:

$$S = \frac{1}{N}\begin{pmatrix} \sum_i (x_i - x_{cm})^2 & \sum_i (x_i - x_{cm})(y_i - y_{cm}) & \sum_i (x_i - x_{cm})(z_i - z_{cm}) \\ \sum_i (x_i - x_{cm})(y_i - y_{cm}) & \sum_i (y_i - y_{cm})^2 & \sum_i (y_i - y_{cm})(z_i - z_{cm}) \\ \sum_i (x_i - x_{cm})(z_i - z_{cm}) & \sum_i (y_i - y_{cm})(z_i - z_{cm}) & \sum_i (z_i - z_{cm})^2 \end{pmatrix}. \tag{8}$$

Transformation to the principal axis system diagonalizes $S$,

$$S = \mathrm{diag}(\lambda_1, \lambda_2, \lambda_3), \tag{9}$$

where the eigenvalues of $S$ are sorted in the descending order $\lambda_1 \geq \lambda_2 \geq \lambda_3$. The first invariant of $S$ gives the squared radius of gyration,

$$\mathrm{Tr}(S) = \lambda_1^2 + \lambda_2^2 + \lambda_3^2 = R_g^2. \tag{10}$$

To measure the deviation from the spherical symmetry, we calculated the asphericity parameter using:

$$\Delta = \frac{3}{2}\frac{\lambda_1^4 + \lambda_2^4 + \lambda_3^4}{(\lambda_1^2 + \lambda_2^2 + \lambda_3^2)^2} - \frac{1}{2}. \tag{11}$$

Here, $\Delta = 0$ corresponds to spherical shape and $\Delta = 1$ denotes a rod-like shape. End-to-end distance of a chain is computed using:

$$\vec{R}_{ee} = \vec{r}_1 - \vec{r}_n, \tag{12}$$

where $r_1$ and $r_n$ are the position of the first and last beads of the RNA chain, respectively.

##### 3.6 Condensate volume

Condensate volume was estimated from the spatial distribution of RNA beads relative to the condensate center of mass (COM). The condensate radius $R$ was defined as the 95th percentile of the resulting distance distribution, corresponding to the radius enclosing 95% of all RNA beads. Assuming an approximately spherical condensate (which is mostly the case, see Fig. 1 in main text), the condensate volume was calculated as $V = \frac{4}{3}\pi R^3$. For statistical analysis, frame-wise volumes were divided into non-overlapping blocks of 100 frames, and the mean volume of each block was used as an independent sample. Reported values correspond to the mean and standard deviation of the block-averaged volumes.

##### 3.7 RNA and $\mathbf{Mg^{2+}}$ concentration of two phases

The RNA concentration inside condensate is computed by averaging the concentrations of individual medium and large condensates:

$$C_{\text{droplet}} = \bar{C}_i, \tag{13}$$

where the concentration of $i$-th condensate is defined as $\bar{C}_i = N_i/V_i$, with $N_i$ denoting the number RNA molecules and $V_i$ the volume of the condensate. The averaging is performed only over condensates containing more than five RNA molecules ($N_i > 5$).

The number of $Mg^{2+}$ ions within the condensate was calculated using the condensate COM as reference. At late stages of the simulation, all 60 RNA chains reside within the condensate. The spatial extent of the condensate was characterized by an effective radius $R$, defined above. An $Mg^{2+}$ ion was considered to be within the condensate if its distance from the condensate COM satisfied $r_{Mg} \leq R$, where $r_{Mg}$ is the COM-to-ion distance. The total number of $Mg^{2+}$ ions within the condensate, $N_{Mg}$, was then obtained by counting all ions satisfying this criterion.

The average $Mg^{2+}$ concentration within the condensate was computed as:

$$C_{Mg} = \left\langle \frac{N_{Mg}}{V} \right\rangle, \tag{14}$$

where $V$ is the condensate volume, and $\langle \cdot \rangle$ denotes the time average over the equilibrated portion of the trajectory. The $Mg^{2+}$ concentration in the dilute phase (outside the condensate) was calculated by subtracting the condensate volume from the total simulation box volume, and the number of $Mg^{2+}$ within the condensate from the total number of $Mg^{2+}$ in the system:

$$C_{Mg}^{dilute} = \left\langle \frac{N_{Mg}^{tot} - N_{Mg}}{V_{box} - V} \right\rangle, \tag{15}$$

where $N_{Mg}^{tot}$ is the total number of $Mg^{2+}$ in the system, $N_{Mg}$ is the number of $Mg^{2+}$ within the condensate, and $V_{box}$ is the simulation box volume.

#### 3.8 Form factor

The form factor is computed using:

$$F(q) = \left\langle \left| \sum_k \exp(i\mathbf{q} \cdot \mathbf{r_k}) \right|^2 \right\rangle_N, \tag{16}$$

where $\mathbf{q}$ is the wave vector, $\mathbf{r}$ is the position of the RNA beads and the averaging is done over the chains at different orientations.

#### 3.9 Stacking propensity

For each RNA chain, each nucleotide was classified as either stacked (S) or unstacked (U) based on its local geometry. Specifically, for nucleotide $i$, we computed the sugar-phosphate dihedral angles $\psi = \psi(p_i, s_i, p_{i+1}, s_{i+1})$ and $\phi_i = \phi(s_i, p_{i+1}, s_{i+1}, p_{i+2})$, and compared them to their reference values $\psi_0$ and $\phi_0$ obtained from the ideal A-form RNA. A nucleotide was considered stacked if both deviations $\delta\psi_i = |\psi_i - \psi_0|$ and $\delta\phi_i = |\phi_i - \phi_0|$ were within 10% of the reference values. Contiguous stacked nucleotides were grouped into stacked segments of length $N_n$, where $n$ indexes each segment and $n_s$ is the total number of such segments in a chain. The stacking propensity $p_s$ was then defined as:

$$p_s = \frac{\sum_n^{n_s} N_n \Theta(N_n - 3)}{N}, \tag{17}$$

where $N$ is the total number of nucleotides in the chain, and $\Theta(x)$ is the Heaviside step function to ensure that only long enough segments contribute to the stacking propensity.

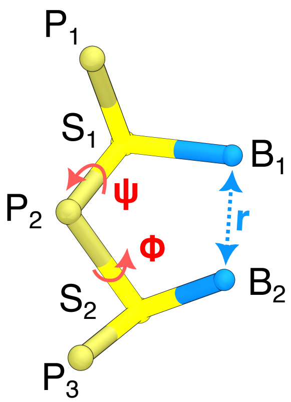

Figure S1: **Structural parameters used to characterize base stacking.** The dihedral angle $\psi$ is defined by the atoms P1–S1–P2–S2, whereas $\phi$ is defined by S1–P2–S2–P3. The parameter $r$ denotes the equilibrium distance between neighboring bases, with values of 4.18 Å for A–A pairs and 4.25 Å for U–U pairs.

#### 3.10 $Mg^{2+}$ binding modes and state assignment

$Mg^{2+}$ binding was classified according to the number and type of phosphate coordination. Bridging $Mg^{2+}$ was first identified by locating $Mg^{2+}$ simultaneously coordinated with at least two phosphates. Each contact was then classified as inner- or outer-sphere using distance-based cutoffs: inner-sphere for $Mg^{2+}$-phosphate distance $< 3.2$ Å, and outer-sphere for those between 3.2 and 6.1 Å. Based on these criteria, bridging configurations were further categorized into three classes: inner-inner (I-I), inner-outer (I-O), and outer-outer (O-O). We also partitioned bridging into inter- and intrachain interactions depending on whether the two phosphates belonged to the same or different RNA chains. For intrachain bridging, adjacent phosphate pairs were excluded to remove trivial local contacts.

To assign a unique coordination state to each $Mg^{2+}$ over time, we constructed a two-dimensional Mg-by-time state matrix. For each frame, a $Mg^{2+}$ ion was marked as occupying one or more bridging classes depending on its coordination environment. Because a single $Mg^{2+}$ can simultaneously satisfy multiple bridging configurations within the same frame, state assignment was encoded using a bitmask representation, where I-I, I-O, and O-O were assigned values of 1, 2, and 4, respectively. These values were combined by bitwise addition to produce composite states corresponding to simultaneous occupancy of multiple coordination modes (e.g., 3 = I-I + I-O, 5 = I-I + O-O, 6 = I-O + O-O, and 7 = I-I + I-O + O-O). $Mg^{2+}$ with no bridging assignment in a given frame were subsequently classified inner-only or outer-only if they appeared in the corresponding single-contact datasets; otherwise, they were labeled as diffusive. This procedure yielded ten distinct $Mg^{2+}$ coordination states: diffusive, I-I, I-O, O-O, I-I + O-O, I-O + O-O, I-I + I-O + O-O, inner, and outer.

#### 3.11 Intra and inter-chain contacts

Intra- and inter-chain contacts were calculated based on the distances between phosphate beads within the same chain or across different chains, respectively. Bead pairs separated by 10 Å or less were considered in contact.

#### 3.12 Radial $Mg^{2+}$ concentration profiles

To characterize the spatial distribution of $Mg^{2+}$ within and surrounding RNA condensates, we calculated the radial $Mg^{2+}$ concentration profiles relative to the condensate COM. For each production trajectory frame, the distance between individual $Mg^{2+}$ ions and the condensate COM were calculated. The radial distance were binned into thin concentric spherical shells with a thickness of ($\Delta r = 0.1$ Å).

The volume of each spherical shell (j), bounded by inner radius ($r_j$) and outer radius

$(r_j + \Delta r)$, was calculated by integrating the spherical volume element:

$$V_j = 4\pi r_j^2 \Delta r. \tag{18}$$

The local $Mg^{2+}$ concentration in each shell was determined by dividing the time-averaged number of $Mg^{2+}$ ions within the shell by the corresponding shell volume:

$$[\mathrm{Mg}^{2+}] = \frac{\langle N_{\mathrm{Mg},j} \rangle}{N_A V_j}, \tag{19}$$

where $\langle N_{\mathrm{Mg},j} \rangle$ is the average number of $Mg^{2+}$ ions in shell $(j)$ across the analyzed simulation trajectory frames, $N_A$ is the Avogadro's constant.

Radial $Mg^{2+}$ concentration profiles were calculated independently for three simulation replicas. The replica-averaged concentration profile was obtained by taking the mean of the three replicas at each radial position. The variability between replica was accessed by the standard deviation.

#### 3.13 Radial Profiles of RNA Chain Conformations

To characterize the spatial variation of RNA conformations within the condensate, we calculated radial profiles of the whole-chain $R_g$ and Flory scaling exponent ($\nu$). For each analyzed simulation frame, the COM of each RNA chain and the condensate COM were determined. The radial position of each chain was defined as the distance between these two COMs. The whole-chain $R_g$ was then assigned to the corresponding radial bin, and the average $R_g$ was calculated over chains and simulation frames contributing to each bin.

The radial Flory scaling exponent was determined using the same spatial assignment. For chains assigned to each radial bin, the mean intrachain distance, $R(s)$, was evaluated as a function of nucleotide sequence separation, $s = |i - j|$. The effective scaling exponent was calculated by fitting the power-law relationship, $R(s) \propto s^{\nu}$, over the selected sequence-separation range. The resulting radial profiles were calculated independently for each simulation replica and subsequently averaged across three independent replicas.

To examine the sensitivity of the $R_g$ profiles to the spatial-assignment procedure, we additionally performed a residue-weighted analysis following Farag et al.[7] In this approach, the whole-chain $R_g$ was assigned to the radial position of each nucleotide relative to the condensate COM. Thus, a single RNA chain could contribute to multiple radial bins, with its contribution to each bin weighted by the number of constituent nucleotide occupying that shell. This differs from the chain-based approach, in which each chain contributes only to the radial bin containing its COM.

#### 3.14 Visualization of $Mg^{2+}$ and Phosphate distributions

To analyze the spatial distribution of phosphate and $Mg^{2+}$ ions, volumetric density maps were generated using the *volmap* tool in VMD. The calculation was performed over the whole production portion of the simulation trajectory. Atomic coordinates were mapped onto a 3D grid with a resolution of 1.0 Å and averaged over all frames. The resulting grid was visualized using the *Volume Slice* representation in VMD. The slice plane was oriented perpendicular to the z-axis.

Table S1: **Summary of condensate simulation conditions and sampling times.** All simulations contained 60 RNA molecules in a cubic simulation box with length 793 Å to fix RNA concentration at 200 $\mu$M. Independent replica simulation times are listed in the same cell and separated by commas.

| Sequence | RNA ($\mu$M) | Na (mM) | Mg (mM) | T (°C) | Simulation time ($\mu$s) | Total ($\mu$s) |
|---|---|---|---|---|---|---|
| $rA_{30}$ | 200 | 300 | 3 | 20 | 4.4, 2.5, 2.0 | 8.9 |
| | | | 3 | 60 | 6.4, 3.0, 4.6 | 13.9 |
| | | | 3 | 100 | 5.0, 1.9, 3.9 | 10.8 |
| | | | 6 | 20 | 4.9, 3.4, 4.1 | 12.4 |
| | | | 6 | 60 | 6.8, 2.8, 3.4 | 13.0 |
| | | | 6 | 100 | 4.3, 2.5, 1.9 | 8.7 |
| | | | 10 | 20 | 4.1, 1.7, 4.9 | 10.7 |
| | | | 10 | 60 | 3.6, 3.3, 1.7 | 8.6 |
| | | | 10 | 100 | 4.1, 1.7, 1.7 | 7.6 |
| | 200 | 50 | 2 | 80 | 3.3 | 3.3 |
| | | | 2 | 90 | 4.4 | 4.4 |
| | | | 2 | 100 | 3,3 | 3.3 |
| | | | 3 | 0 | 2.6 | 2.6 |
| | | | 3 | 20 | 2.5 | 2.5 |
| | | | 3 | 40 | 2.6 | 2.6 |
| | | | 3 | 60 | 2.5 | 2.5 |
| | | | 3 | 80 | 2.2 | 2.2 |
| | | | 3 | 100 | 4.8 | 4.8 |
| $rU_{30}$ | 200 | 300 | 3 | 20 | 3.2, 3.0, 2.1 | 8.3 |
| | | | 3 | 60 | 4.0, 3.3, 2.2 | 9.6 |
| | | | 3 | 100 | 4.1, 3.0, 2.0 | 9.1 |
| | | | 6 | 20 | 3.1, 2.7, 3.1 | 8.9 |
| | | | 6 | 60 | 3.1, 4.6, 3.0 | 10.8 |
| | | | 6 | 100 | 2.4, 4.9, 1.9 | 9.1 |
| | | | 10 | 20 | 2.8, 2.3, 1.7 | 6.7 |
| | | | 10 | 60 | 2.6, 2.3, 1.6 | 6.5 |
| | | | 10 | 100 | 2.9, 4.4, 2.9 | 10.3 |

Table S2: **Dilute vs. Dense $Mg^{2+}$ concentrations.** $Mg^{2+}$ concentrations in the two phases at different temperatures and input concentrations. The background $Na^{+}$ concentration is 300 mM.

| **Input (mM)** | **T (°C)** | **Dilute (mM)** | **Dense (M)** |
|---|---|---|---|
| 3 | 0 | 0.43 | 1.93 |
| | 20 | 0.36 | 2.06 |
| | 60 | 0.27 | 2.26 |
| | 100 | 0.17 | 2.48 |
| 6 | 20 | 3.01 | 2.32 |
| | 60 | 2.90 | 2.54 |
| | 100 | 2.74 | 2.81 |
| 10 | 20 | 6.76 | 2.54 |
| | 60 | 6.62 | 2.84 |
| | 100 | 6.54 | 2.96 |

Table S3: **Summary of $rA_{30}$ condensate properties.** All simulations contained 60 $rA_{30}$ RNA chains in a cubic simulation box with a side length of 793 Å, corresponding to a nominal RNA concentration of 200 $\mu$M. Condensate radii, volumes, and RNA concentration values are reported as mean ± standard deviation across three independent replica (n = 3). When the standard deviation is zero at the reported precision, only the mean is shown.

| RNA | Mg (mM) | T (°C) | Radius (Å) | Volume (nm$^3$) | [RNA] (mM) |
|---|---|---|---|---|---|
| | 3 | 20 | 55.8 | 727.8 | 136.9 |
| | 3 | 60 | 54.9 | 693.1 | 143.7 |
| | 3 | 100 | 53.9 ± 0.1 | 657 ± 2 | 151.6 ± 0.1 |
| | 6 | 20 | 56.1 ± 0.1 | 741 ± 2 | 134.5 |
| $rA_{30}$ | 6 | 60 | 54.8 ± 0.5 | 690 ± 20 | 144.3 ± 0.4 |
| | 6 | 100 | 54.1 ± 0.3 | 660 ± 10 | 150.5 ± 0.2 |
| | 10 | 20 | 55.8 ± 0.2 | 726 ± 8 | 137.2 ± 0.2 |
| | 10 | 60 | 54.7 ± 0.1 | 687 ± 4 | 145.1 ± 0.1 |
| | 10 | 100 | 54.1 ± 0.4 | 660 ± 10 | 150.2 ± 0.3 |

Table S4: Same as Table S3, but for $rU_{30}$.

| RNA | Mg (mM) | T (°C) | Radius (Å) | Volume (nm$^3$) | [RNA] (mM) |
|---|---|---|---|---|---|
| | 3 | 20 | 57.9 | 813.1 | 122.5 |
| | 3 | 60 | 56.8 ± 0.1 | 769 ± 2 | 129.6 ± 0.4 |
| | 3 | 100 | 55.6 ± 0.1 | 721 ± 2 | 138.1 ± 0.4 |
| | 6 | 20 | 57.5 ± 0.2 | 800 ± 10 | 125 ± 2 |
| $rU_{30}$ | 6 | 60 | 56.6 ± 0.3 | 760 ± 10 | 131 ± 2 |
| | 6 | 100 | 55.6 ± 0.6 | 720 ± 20 | 139 ± 4 |
| | 10 | 20 | 57.5 ± 0.4 | 800 ± 20 | 125 ± 2 |
| | 10 | 60 | 56.2 ± 0.2 | 742 ± 6 | 134 ± 1 |
| | 10 | 100 | 55.5 ± 0.7 | 720 ± 30 | 139 ± 5 |

### 4 Effect of monovalent ions on phase separation

Consistent with some experiments,[8–10] we find that the phase boundary exhibits a compensatory dependence on monovalent and divalent ion concentrations, such that decreasing monovalent ion concentrations shifts the phase boundary to higher $Mg^{2+}$ concentrations. This behavior is consistent with mean-field electrostatic models, including Debye-Hückel screening and Manning's counterion condensation theory.[11,12] In this regime, $Na^{+}$ and $Mg^{2+}$ act as partially interchangeable contributors to electrostatic screening, leading to a trade-off between the two. However, other experimental studies have reported more complex, sequence-dependent behaviors, where monovalent ions can compete with or suppress $Mg^{2+}$-driven condensation, highlighting the role of specific ion-RNA interactions beyond mean-field electrostatic dependence.[13,14] Similar dependent behaviors have also been observed in coacervate and other condensate systems.[15–18]

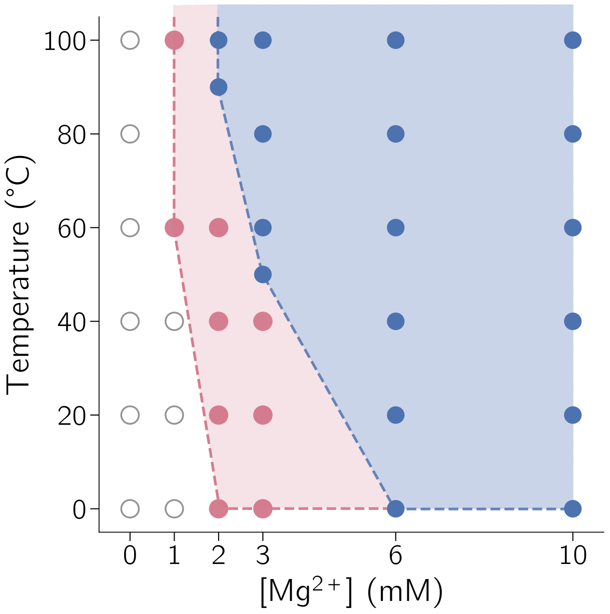

Figure S2: Phase diagram of $rA_{30}$, showing that phase separation occurs upon heating, consistent with LCST-type behavior. Increasing $Mg^{2+}$ concentration promotes phase separation while decreasing $Na^{+}$ concentration decreases phase separation propensity.

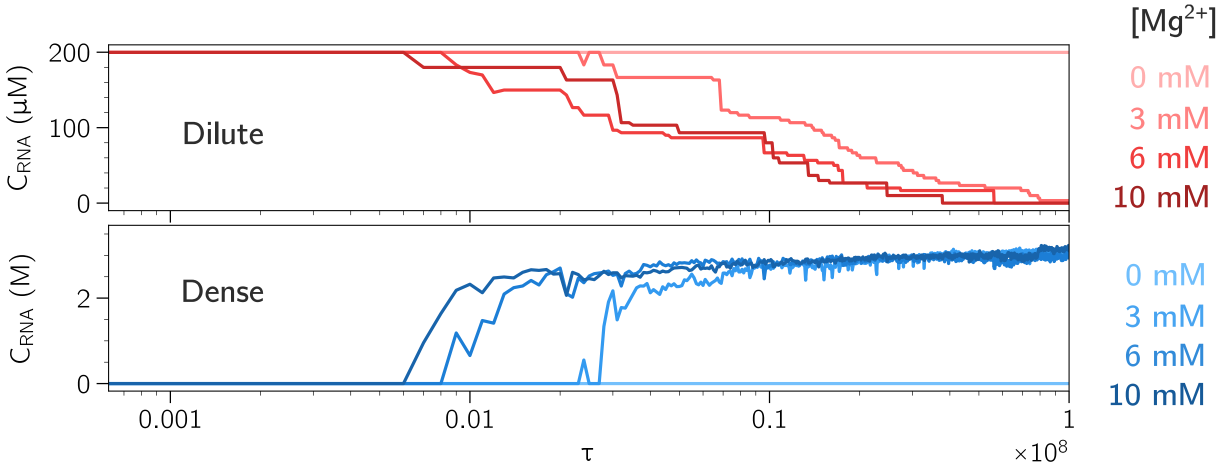

Figure S3: **$Mg^{2+}$-induced phase separation of $rA_{30}$.** RNA concentration in the dilute (gray) and dense (red) phases at T = 20 °C in a mixture of 300 mM $Na^{+}$ and $[Mg^{2+}]$ = 0, 3, 6 and 10 mM.

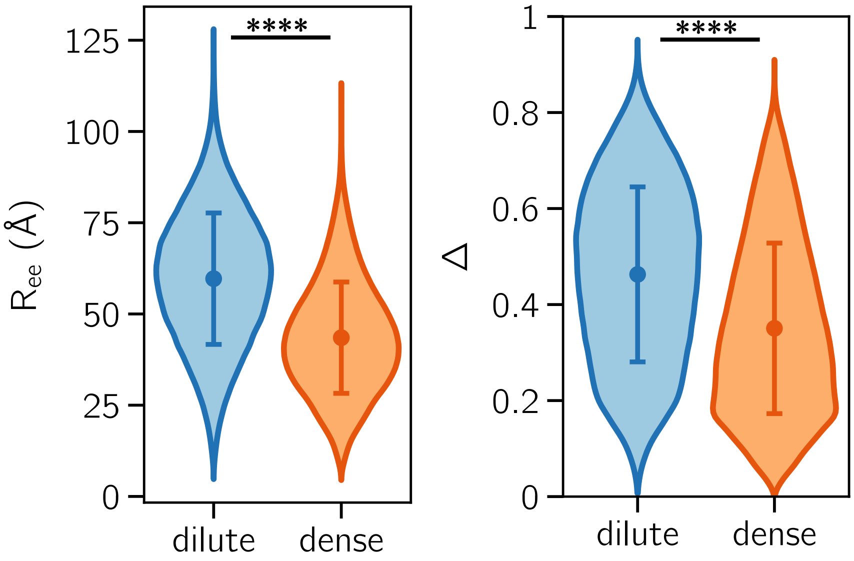

Figure S4: **Size and shape of ssRNA within droplet**. Left and right: end-to-end distance ($R_{ee}$) and asphericity ($\Delta$), respectively. Asterisks indicate statistical significance ($**** : p < 0.0001$).

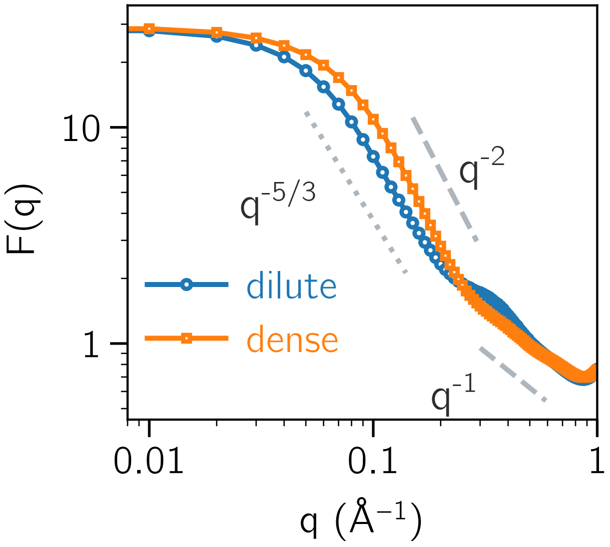

Figure S5: Form factors in the dilute and dense phases. Reference lines with $q^{-5/3}$, $q^{-2}$, and $q^{-1}$ indicate scaling expected for self-avoiding walk (SAW), Gaussian, and rod-like conformations, respectively.

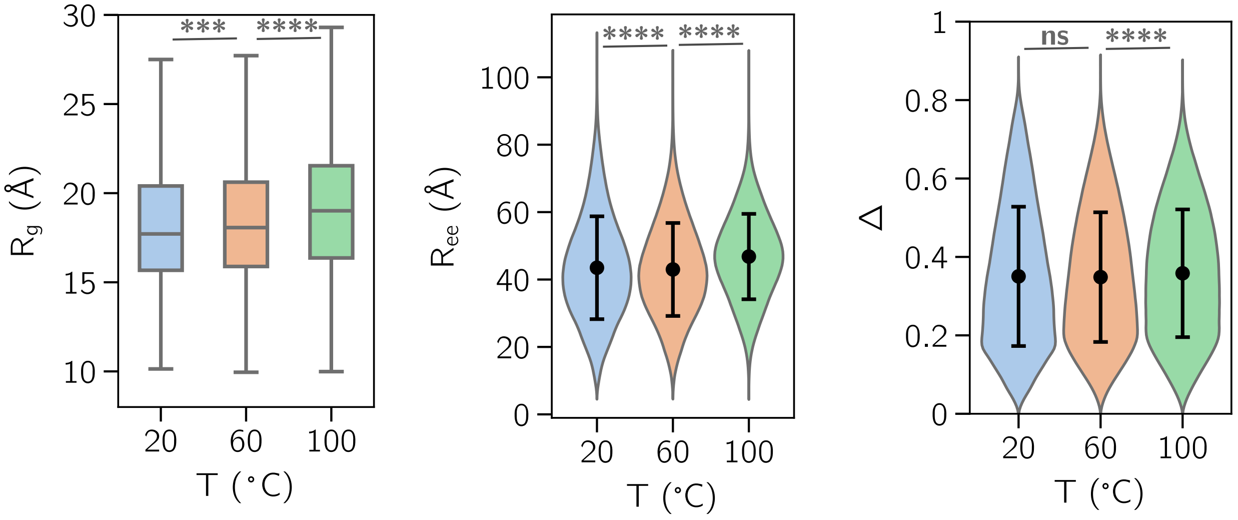

Figure S6: Distribution of (left) $R_g$ (middle) $R_{ee}$ and (right) $\Delta$ for $rA_{30}$ in condensate. Asterisks indicate statistical significance (****: $p < 0.0001$; ***: $p < 0.001$; ns: not significant).

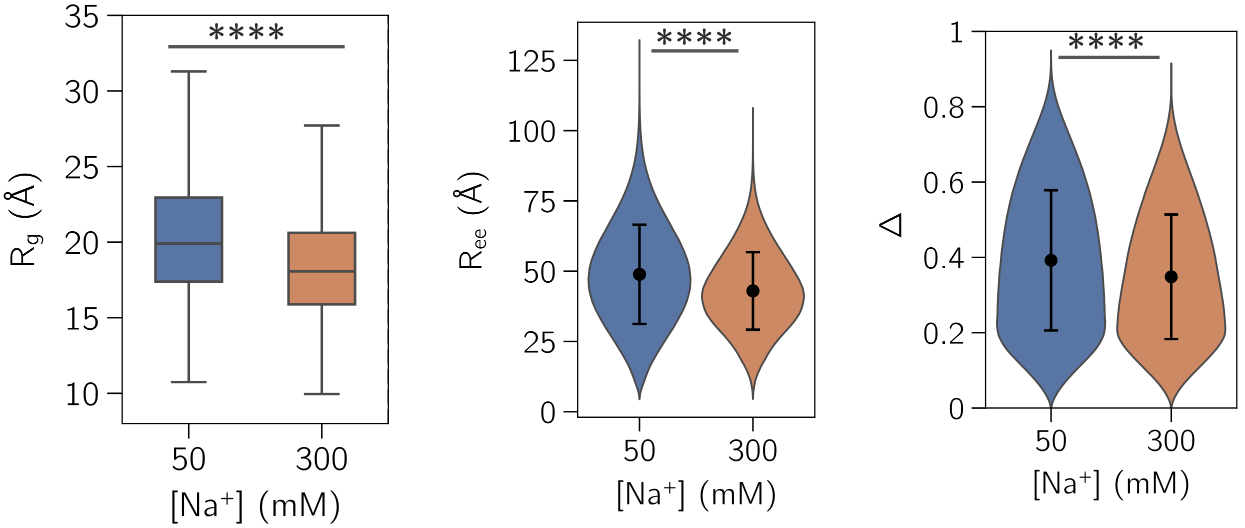

Figure S7: Same as Fig. S6, but at different monovalent ion concentrations.

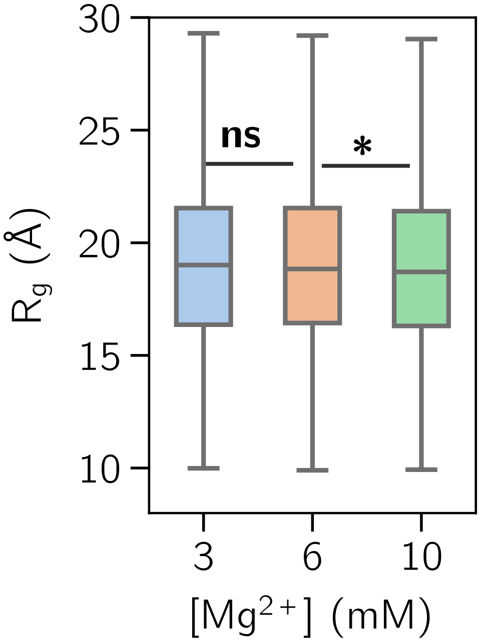

Figure S8: Distribution of $R_g$ for $rA_{30}$ at different $Mg^{2+}$ concentration. Asterisks indicate statistical significance ($*$ : p $<$ 0.05; ns : not significant).

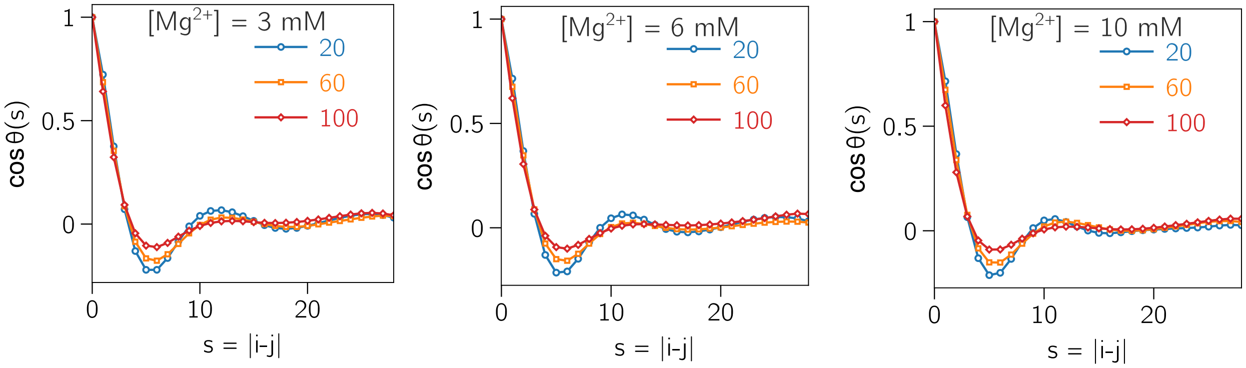

Figure S9: $\cos\theta(s)$ vs. $s$ in $rA_{30}$ condensate at different temperatures and $Mg^{2+}$ concentrations.

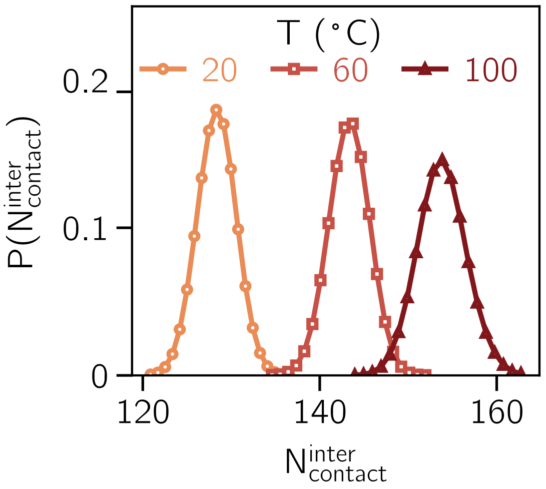

Figure S10: Number of interchain contacts per RNA chain within condensate.

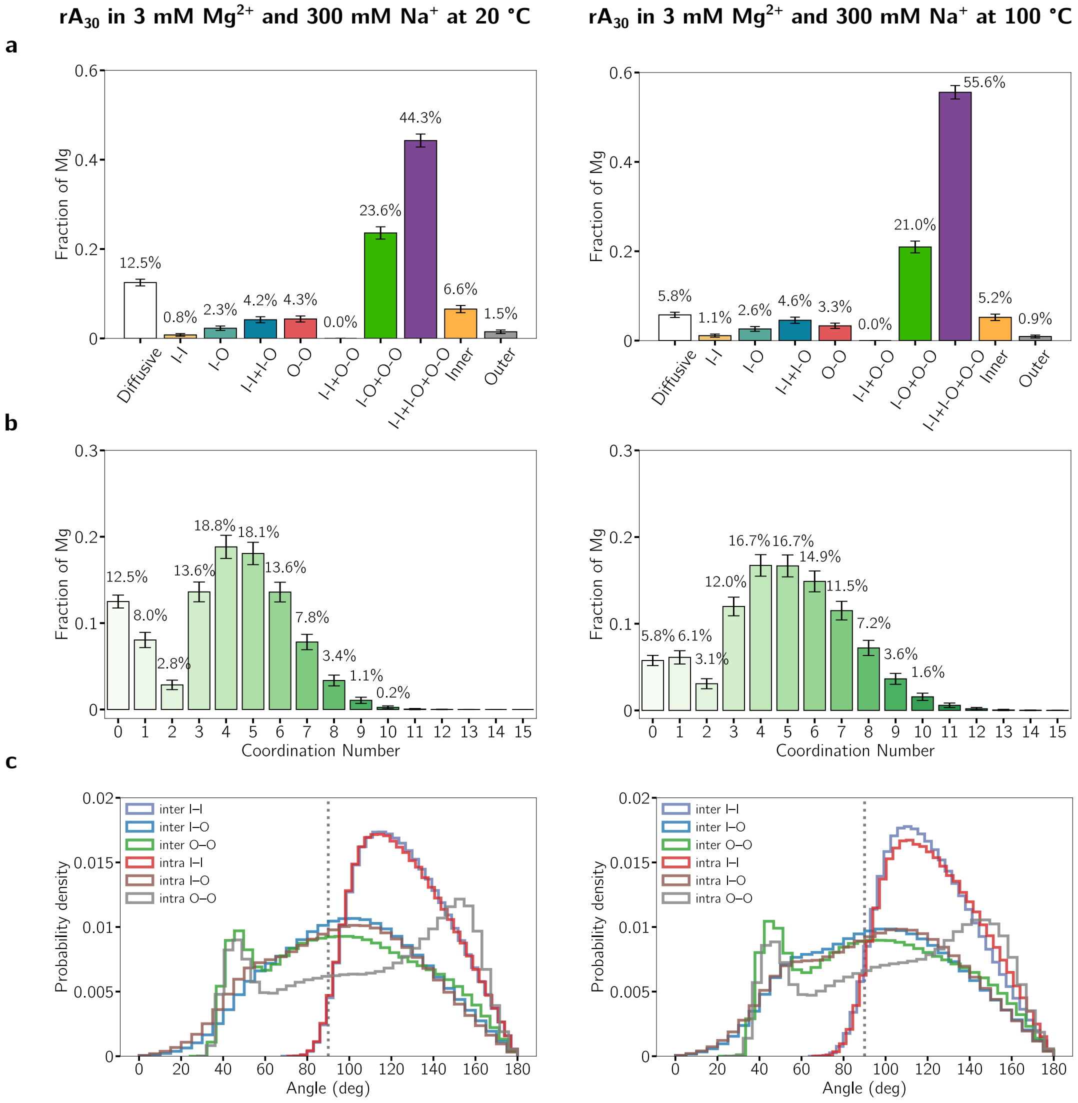

Figure S11: **(a)** Population of $Mg^{2+}$ binding modes in $rA_{30}$ condensates at 3 mM $Mg^{2+}$ and 300 mM $Na^{+}$ at 20 °C (left) and 100 °C(right). Error bars represent the standard deviation across trajectory blocks. **(b)** Distribution of $Mg^{2+}$ coordination numbers under the same conditions. Increasing temperature shifts the coordination distribution toward higher coordination numbers. **(c)** Angles of inter- and intrachain $Mg^{2+}$ bridging geometries for inner-sphere (I-I), mixed inner/outer-sphere (I-O), and outer-sphere (O-O) coordination states remain largely unchanged with temperature. Intrachain bridging is predominantly restricted to angles greater than 80° due to steric constraints imposed by chain connectivity, whereas interchain bridges exhibit substantially broader angular distributions.

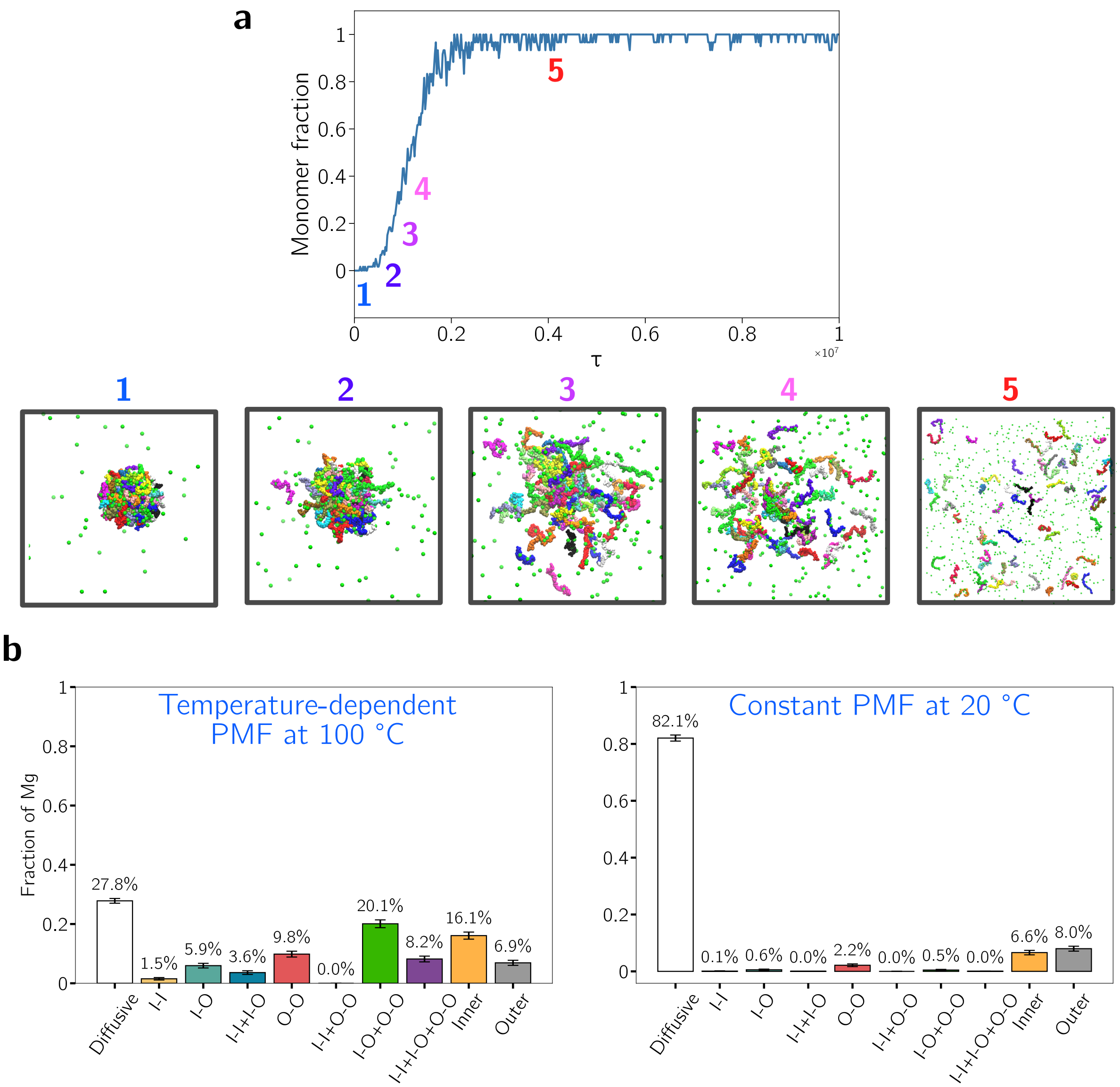

Figure S12: **Effect of temperature-dependent $Mg^{2+}$-P interactions on RNA condensate stability and interchain bridging.** **(a)** Control simulation performed at 100 °C in the presence of 50 mM $Na^{+}$ and 3 mM $Mg^{2+}$, in which $Mg^{2+}$–P PMF was held at 20 °C. Starting from the preformed condensate obtained at 100 °C (snapshot 1), the RNA chains progressively disperse (snapshots 2 to 5), as indicated by the increasing monomer fraction. **(b)** Distribution of $Mg^{2+}$ coordination in the original simulation using the temperature-dependent PMF at 100 °C (left) and the control simulation using 20 °C PMF at 100 °C (right). The fraction of $Mg^{2+}$ ions in interchain bridging decreases from 49.1% to 3.1% upon replacing the temperature-dependent PMF, accompanying the dissolution of the condensate.

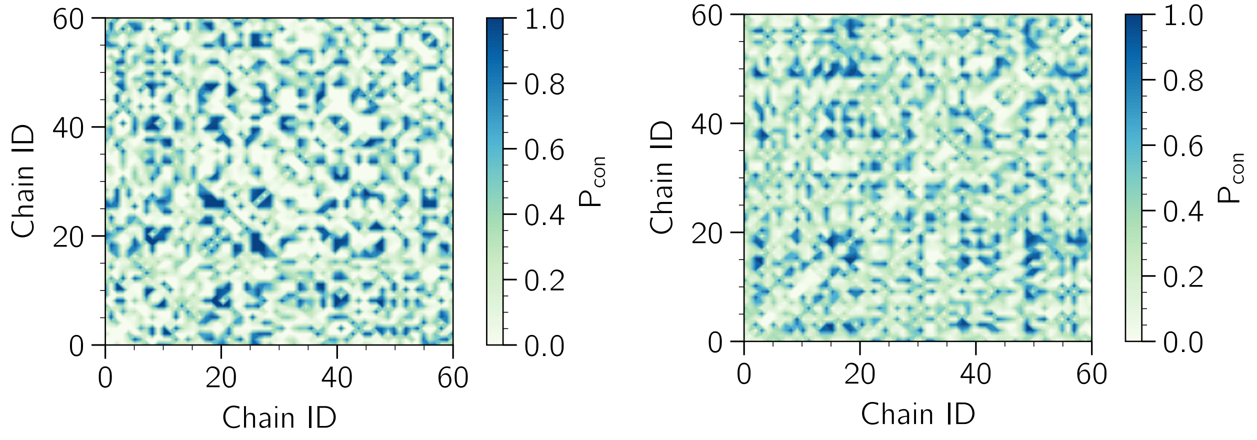

Figure S13: Contact map between chains in the droplet for (left) $rA_{30}$ and (right) $rU_{30}$.

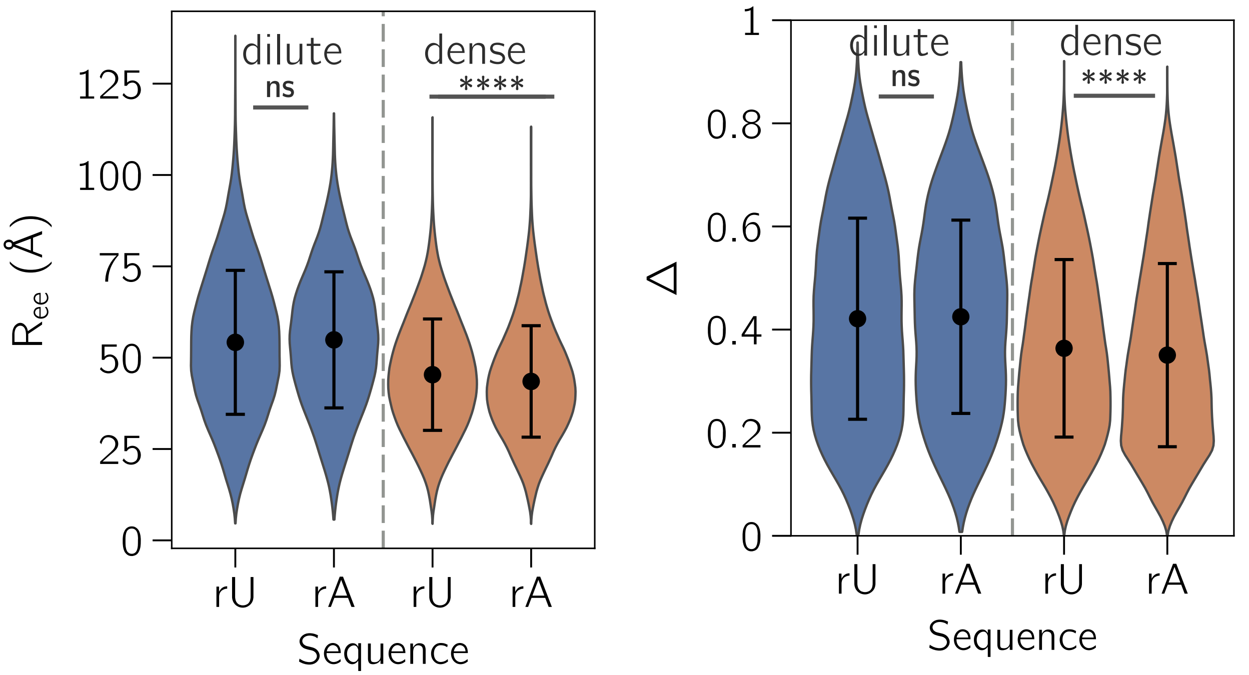

Figure S14: Distribution of $R_{ee}$ (left) and $\Delta$ (right) for $rA_{30}$ and $rU_{30}$ in dilute and dense phases. Asterisks indicate statistical significance (**** : $p < 0.0001$; ns : not significant).

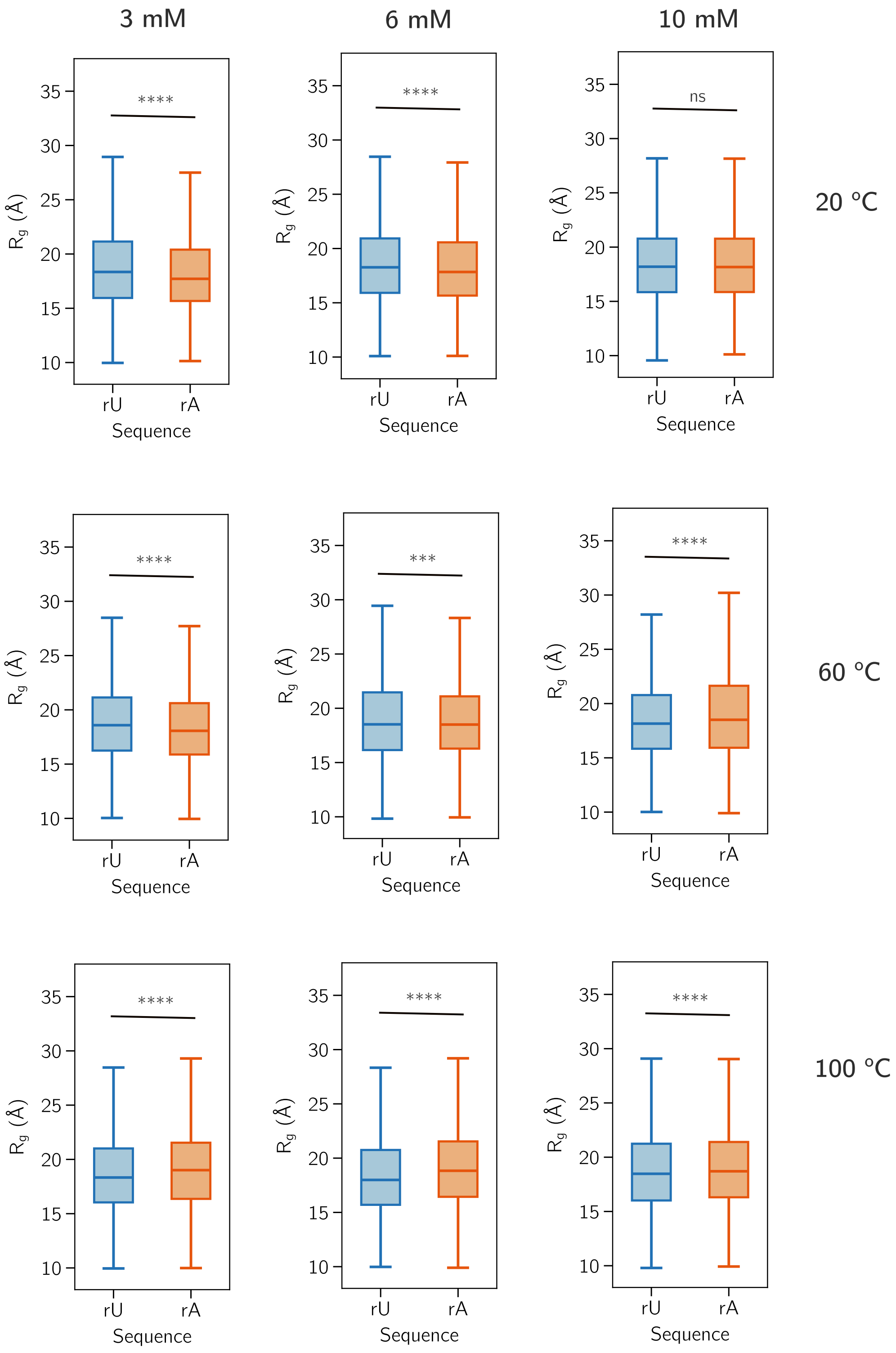

Figure S15: Distribution of $R_g$ for $rA_{30}$ and $rU_{30}$ in condensate at different conditions. Asterisks indicate statistical significance (*** : p < 0.001, **** : p < 0.0001; ns : not significant).

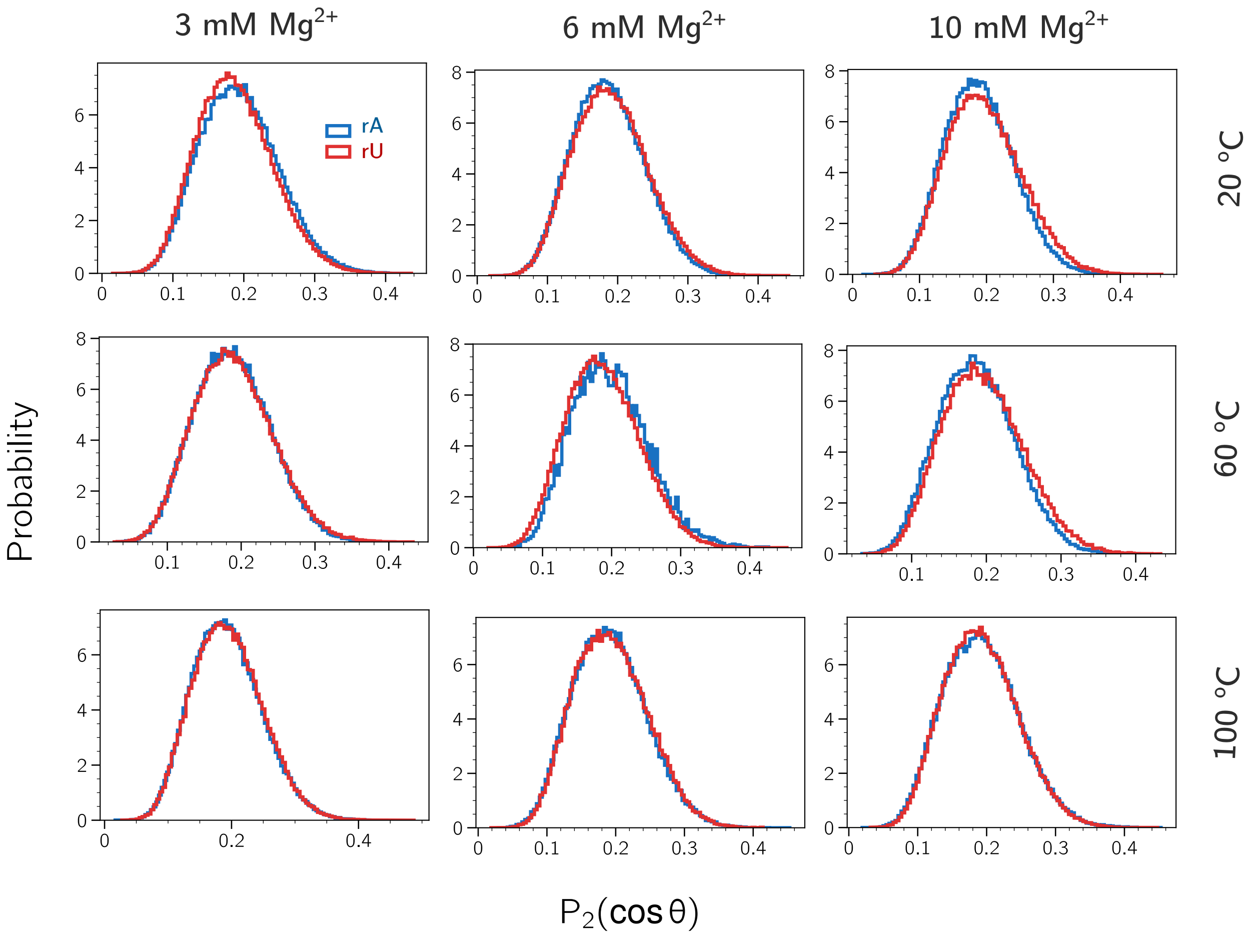

Figure S16: Nematic order parameter of chains within $rA_{30}$ and $rU_{30}$ condensates at different conditions.

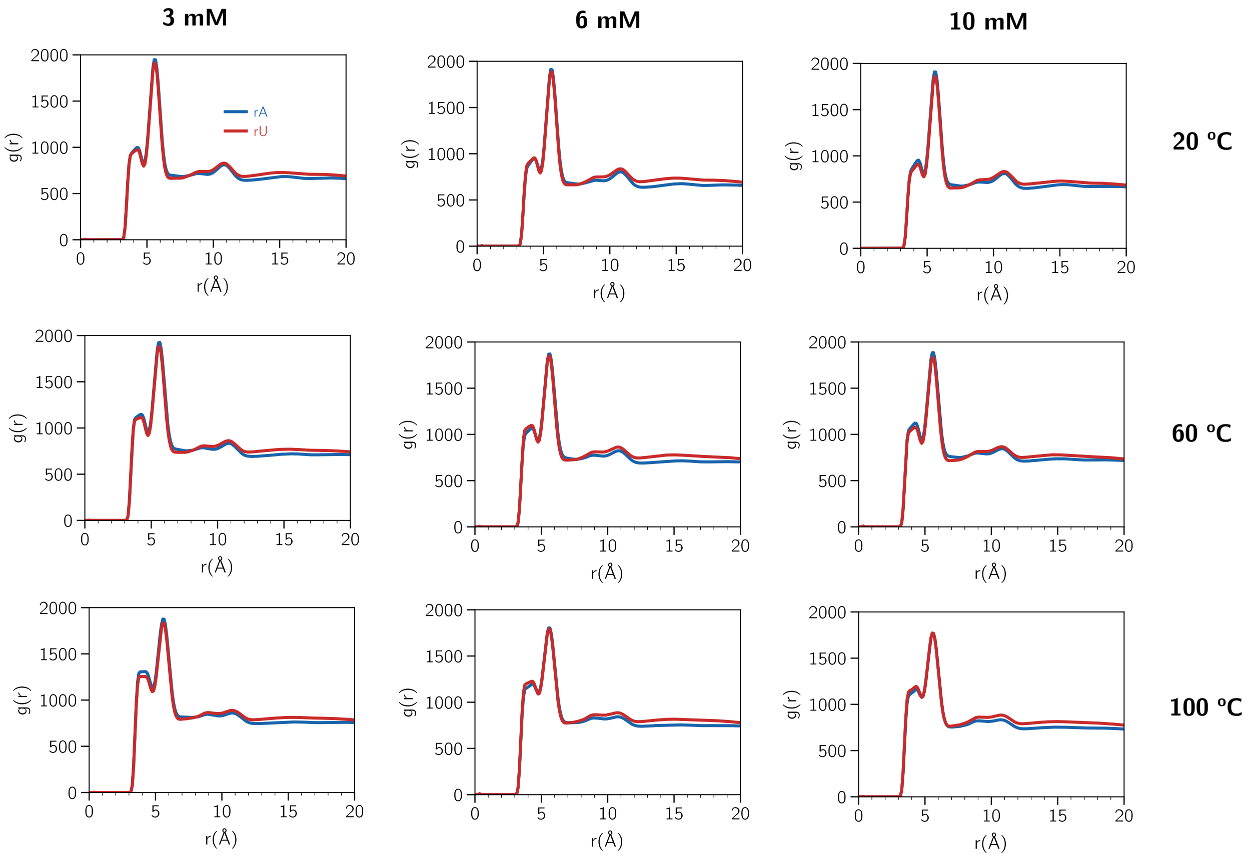

Figure S17: Radial distribution function between RNAs in condensate at different conditions.

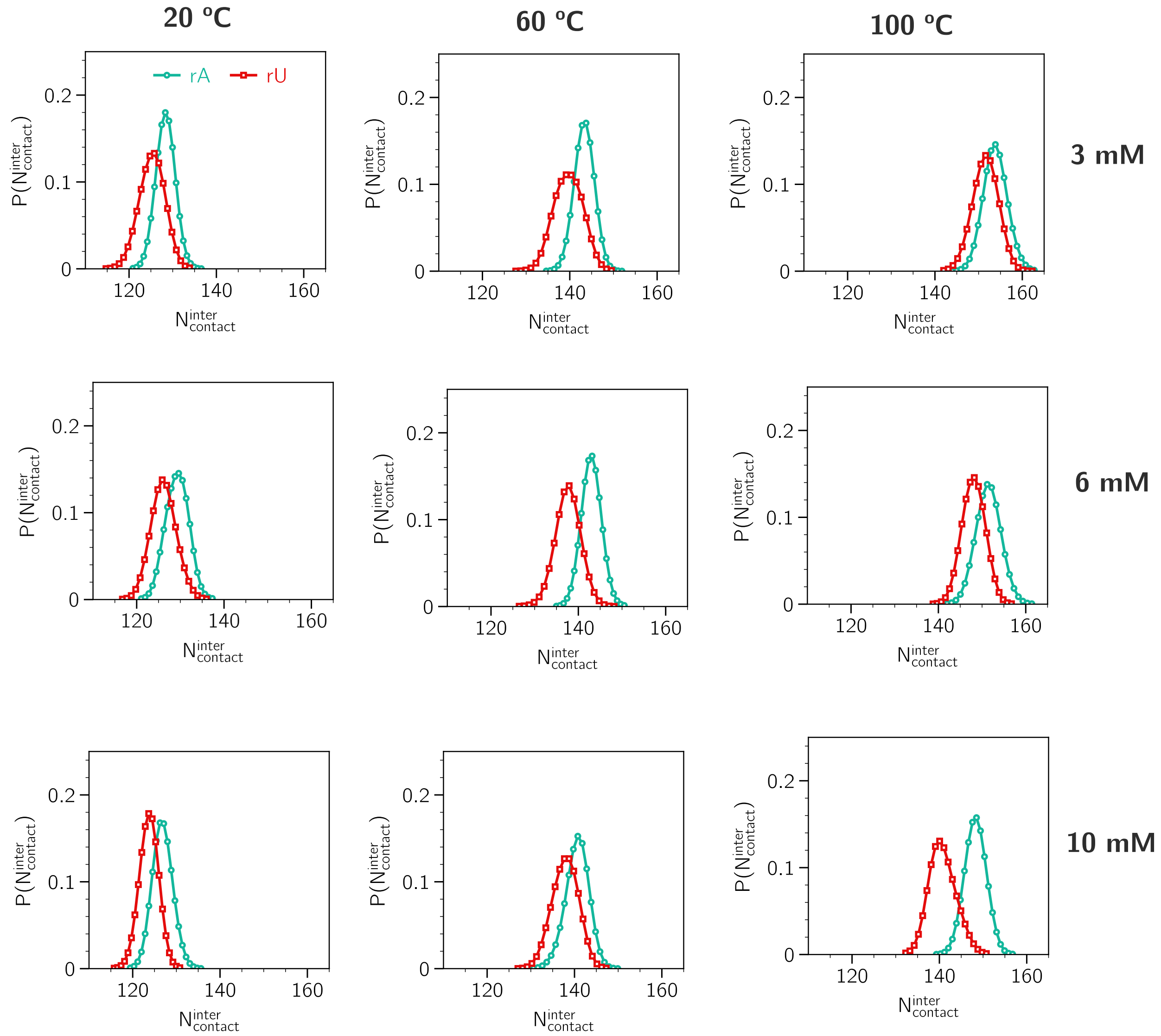

Figure S18: Number of interchain contact per chain in the condensate at different conditions.

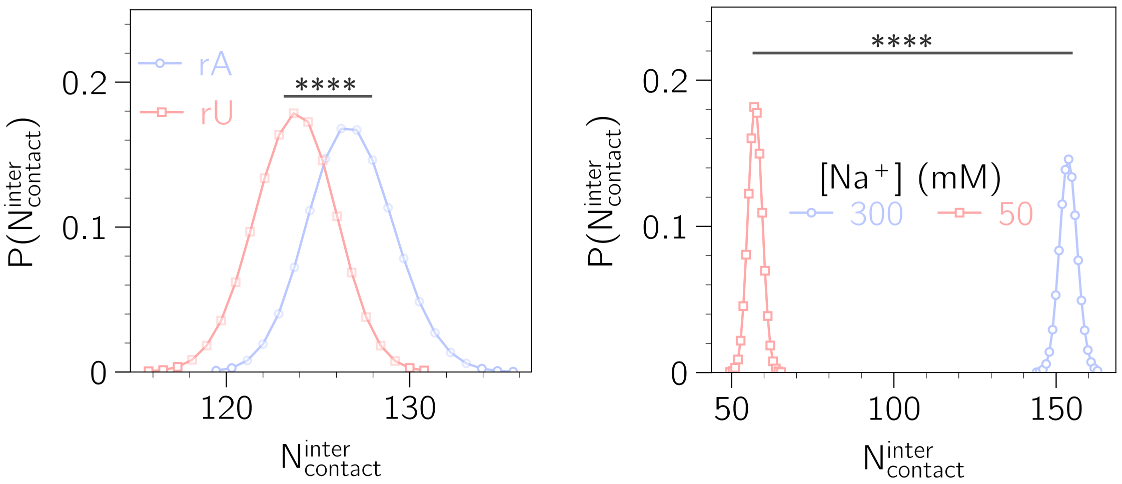

Figure S19: Distribution of interchain contact in condensate (Left) for $rA_{30}$ vs. $rU_{30}$, and (Right) 50 vs. 300 mM $Na^+$. Asterisks indicate statistical significance (****: $p < 0.0001$).

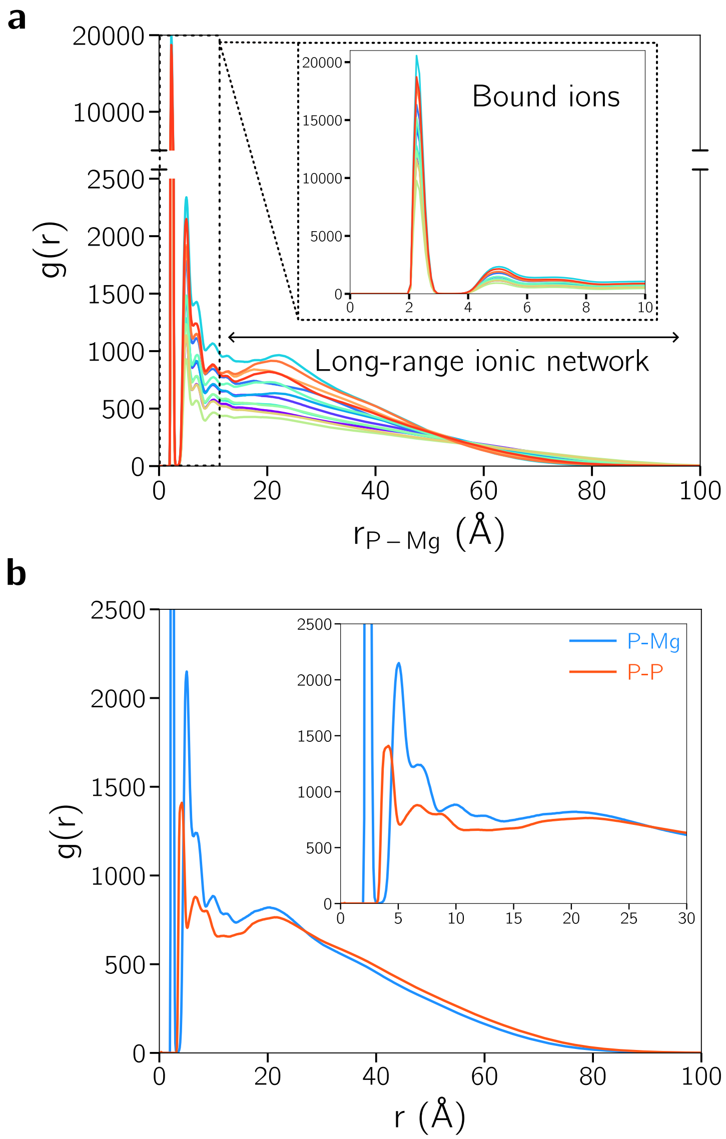

Figure S20: **(a)** Radial distribution functions between phosphate beads and $Mg^{2+}$, revealing a heterogeneous ionic environment. Insets highlight short-range correlations. **(b)** Comparison of intra- vs. interchain correlations.

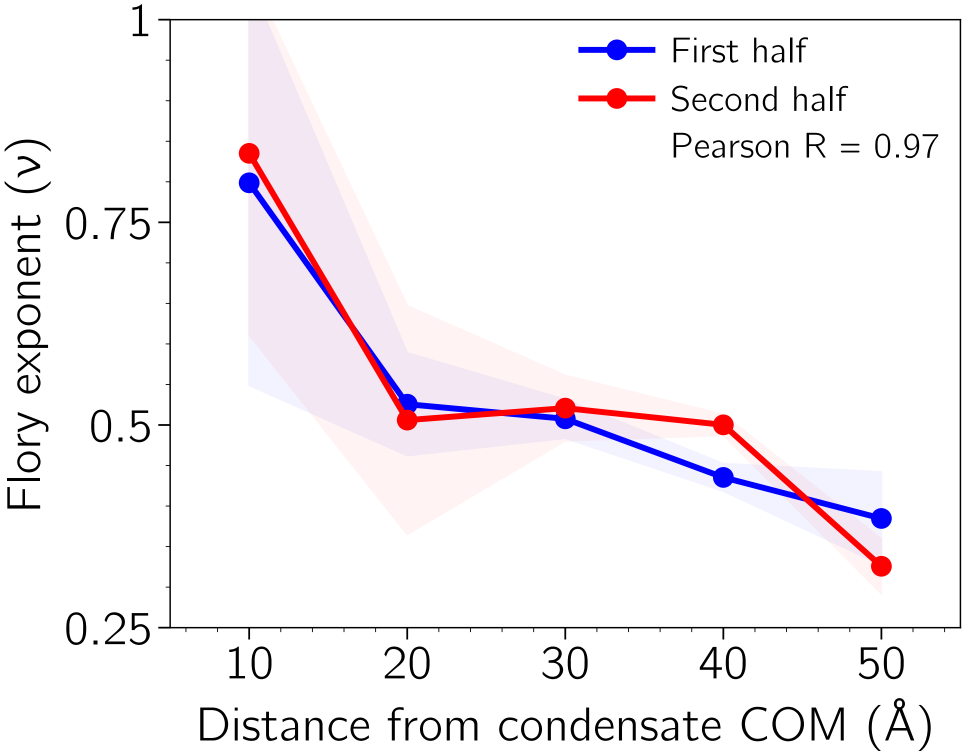

Figure S21: **Convergence of the Flory exponent as a function of radial distance.** Radial scaling-exponent profiles were calculated independently from the first (in blue) and the second (in red) non-overlapping halves of the production trajectories. Solid lines represent the replica-averaged values, and the shaded region represent ±1 standard error of the mean across three independent replicas. The Pearson correlation coefficient between the two profiles was R = 0.97.

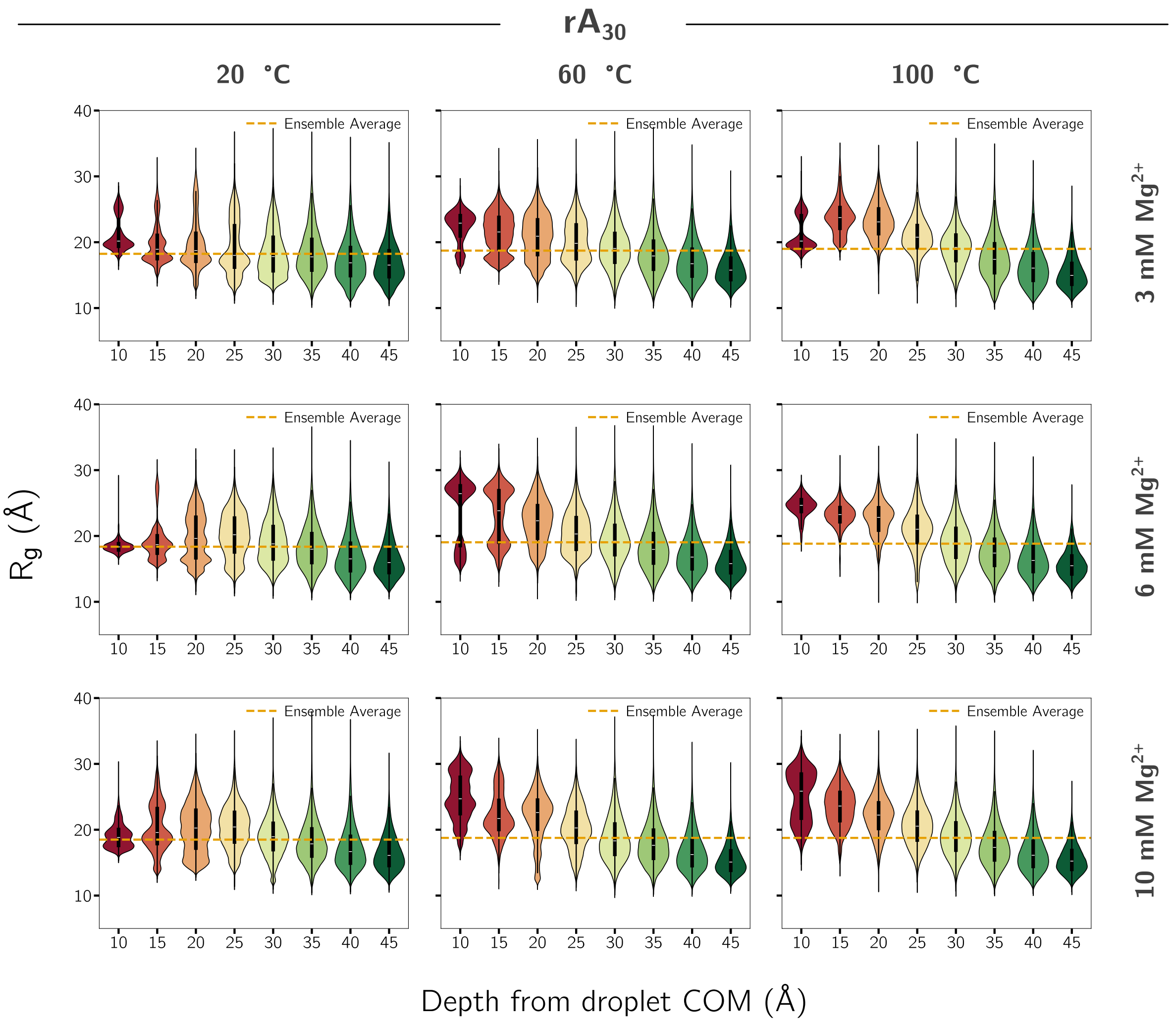

Figure S22: $R_g$ of individual chains in $rA_{30}$ condensate at 300 mM $Na^+$ and various $Mg^{2+}$ concentrations as a function of radial distance.

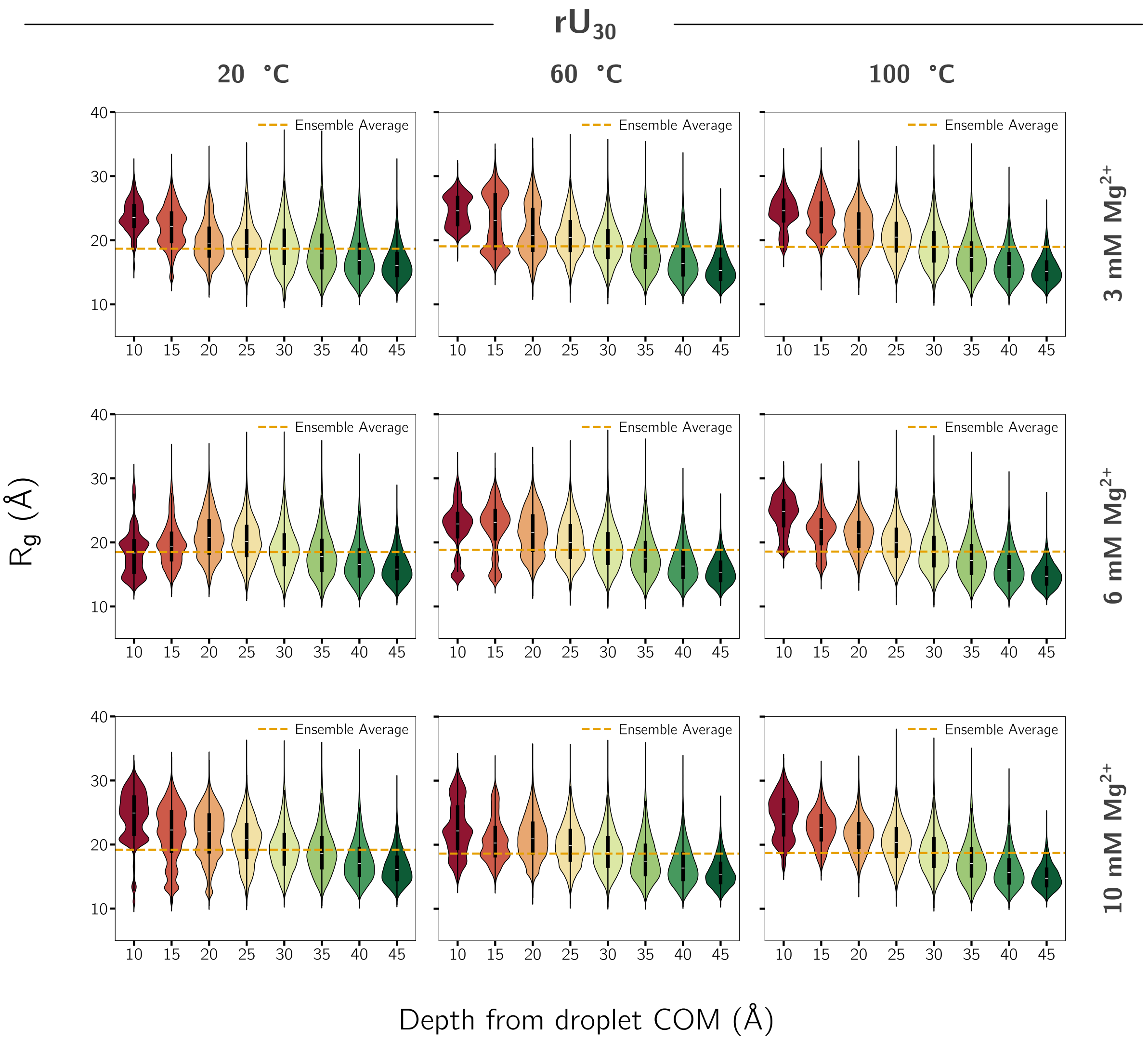

Figure S23: Same as Fig. S22, but for $rU_{30}$.

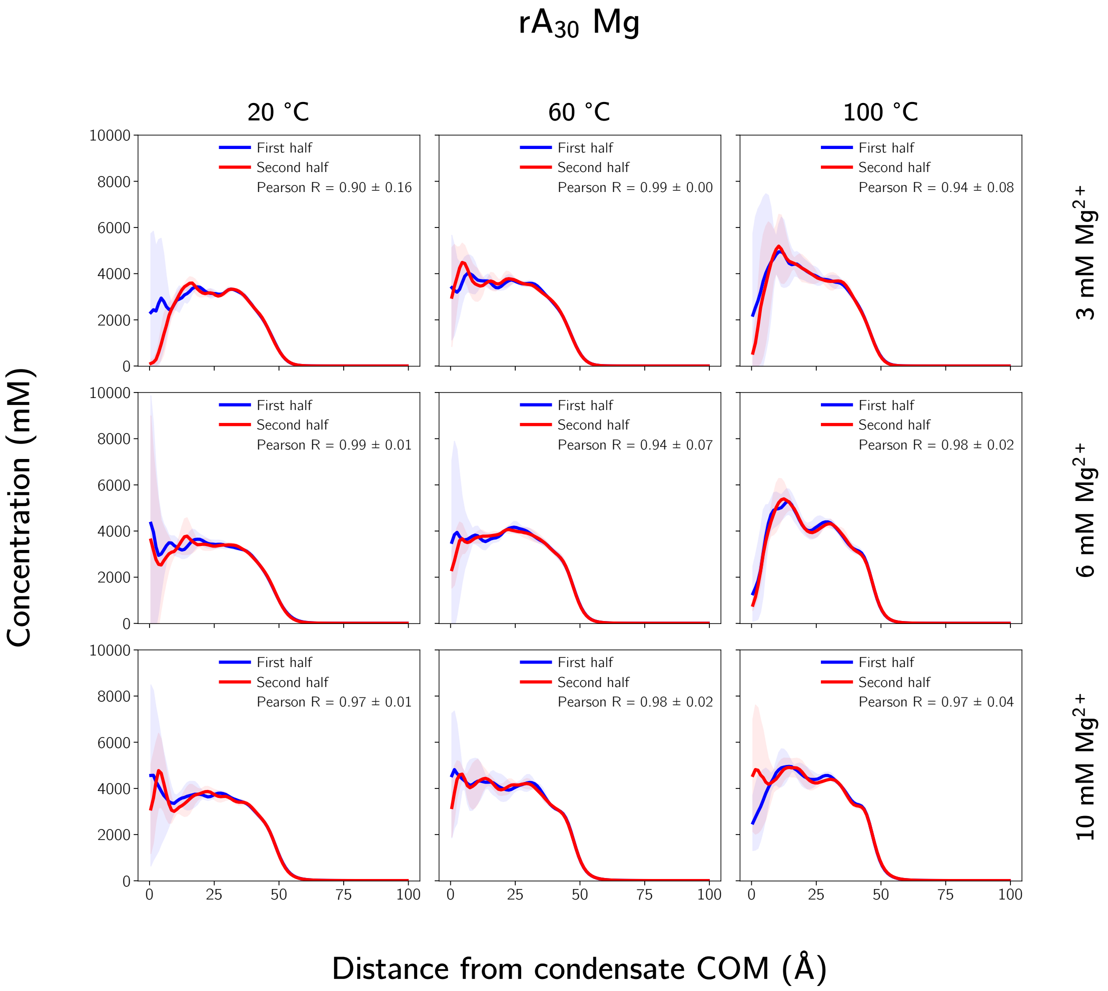

Figure S24: **Temporal convergence of radial $Mg^{2+}$ concentration profiles in $rA_{30}$ condensates.** Radial $Mg^{2+}$ concentration profiles calculated from the first (blue) and second (red) non-overlapping halves of the production trajectories. Solid lines are means across three independent replicas, and the shaded regions are ±1 standard deviation. Pearson correlation coefficient were calculated for each replica and are reported as mean ± SD.

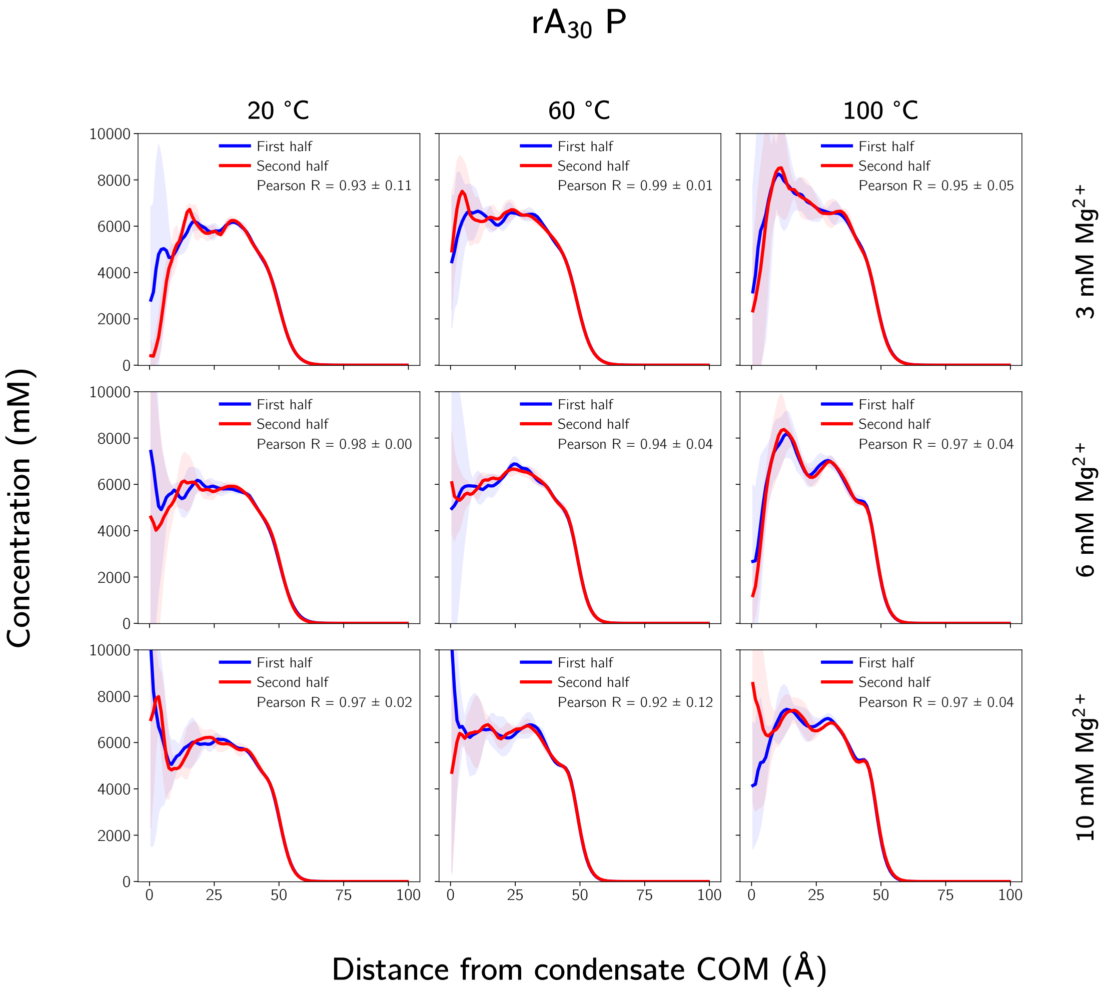

Figure S25: Same as Fig. S24 but for $rA_{30}$ phosphates.

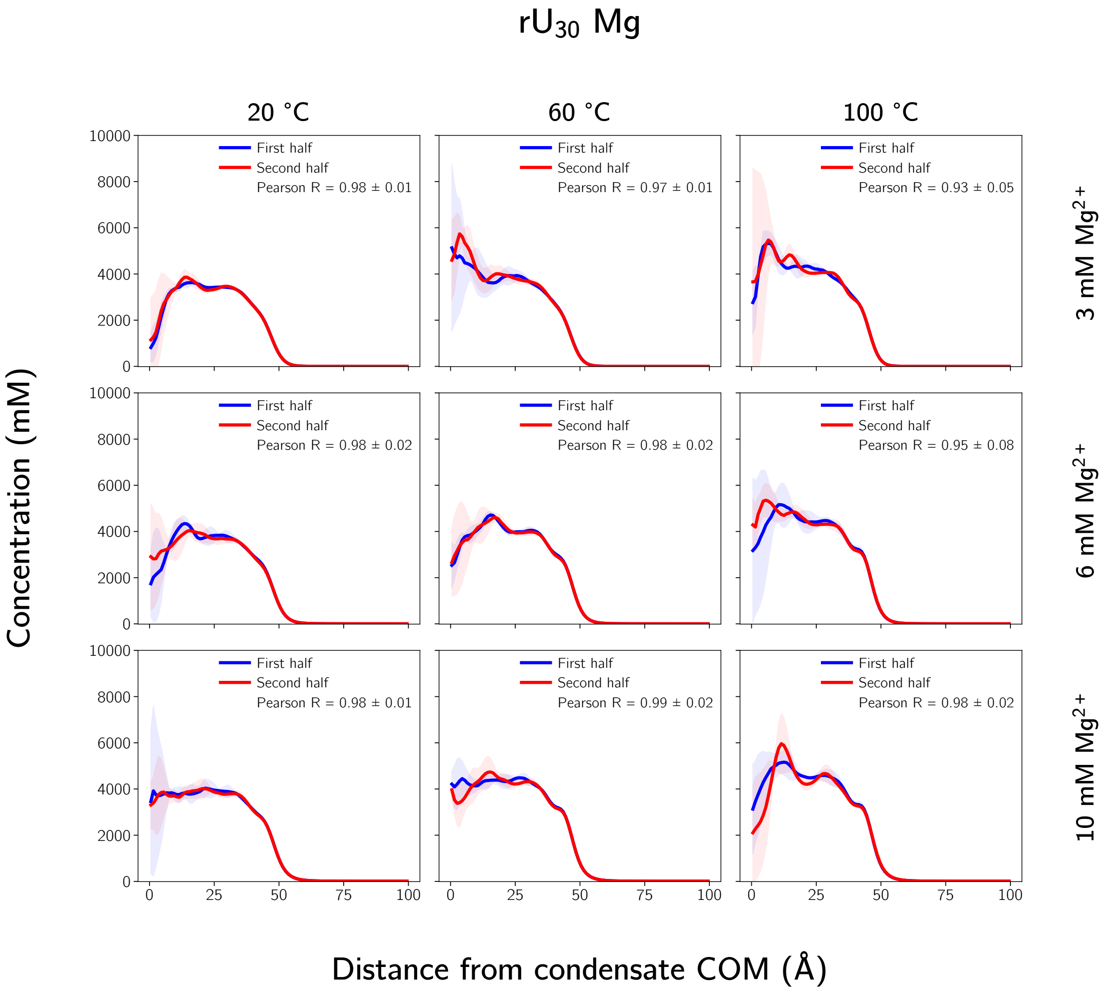

Figure S26: Same as Fig. S24 but for $rU_{30}$ $Mg^{2+}$.

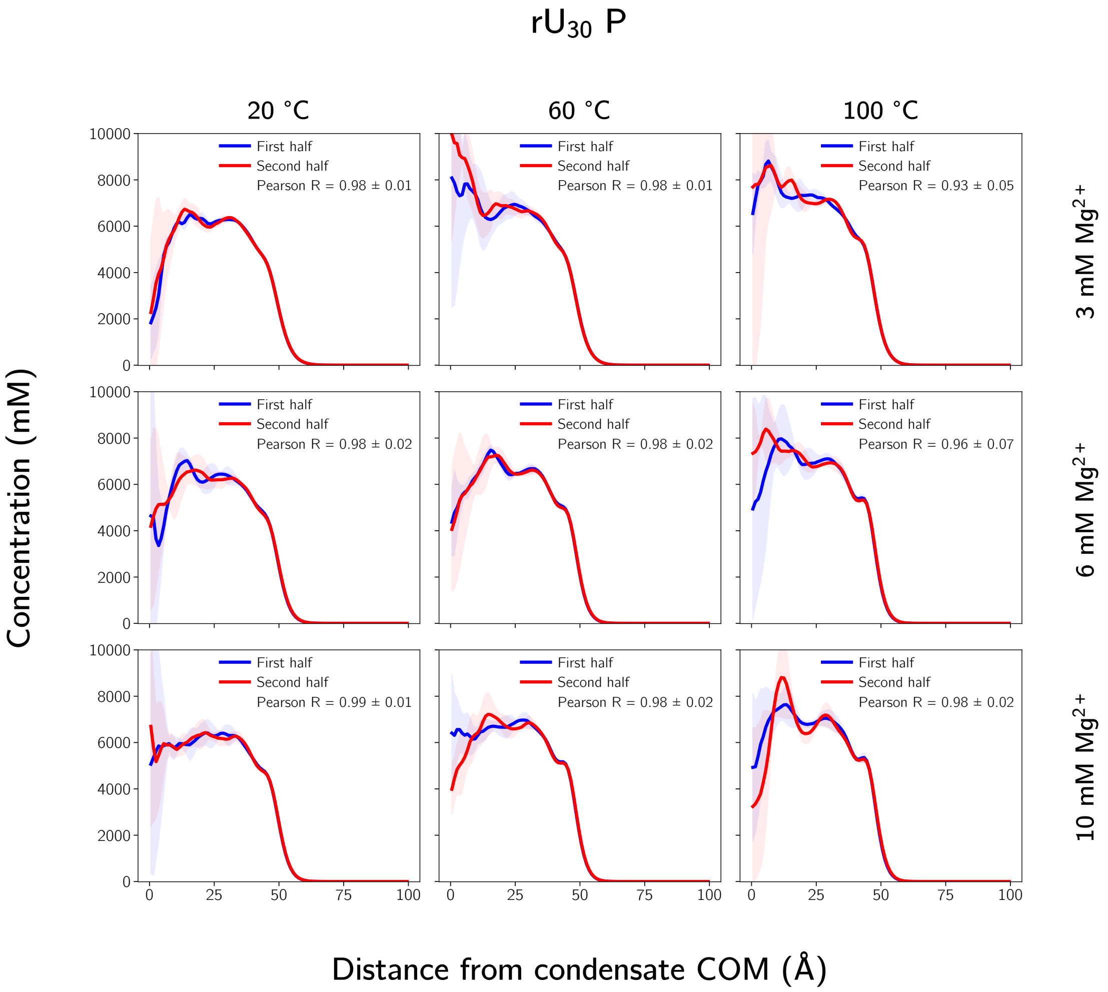

Figure S27: Same as Fig. S24 but for $rU_{30}$ phosphates.

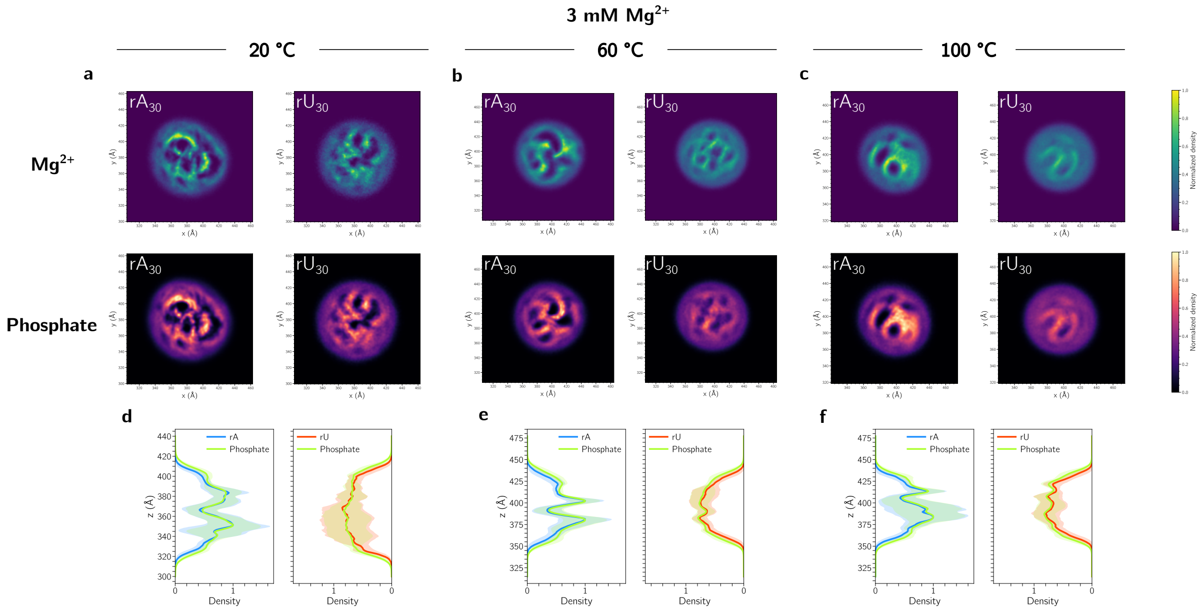

Figure S28: **a-c.** Representative slices along the z-axis of $rA_{30}$ and $rU_{30}$ condensates at 20, 60, and 100 °C in 3 mM $Mg^{2+}$. **d-f.** One-dimensional $Mg^{2+}$ density profiles along the z-axis obtained from cylindrical slices with radius of 10 Å at same conditions.

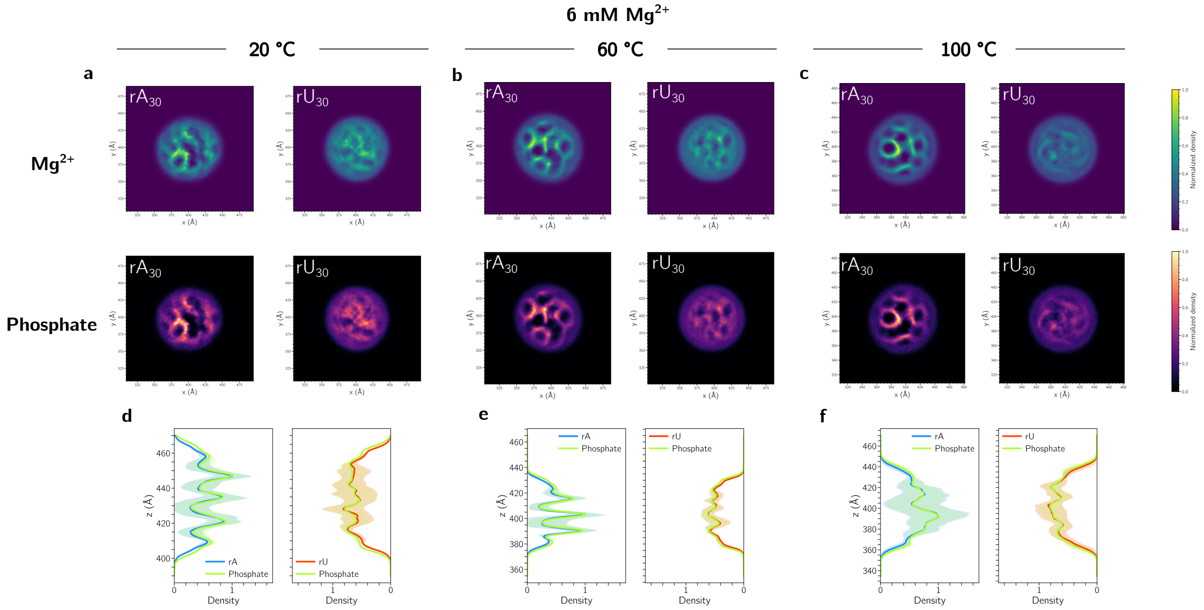

Figure S29: Same as Fig. S28, but in 6 mM $Mg^{2+}$.

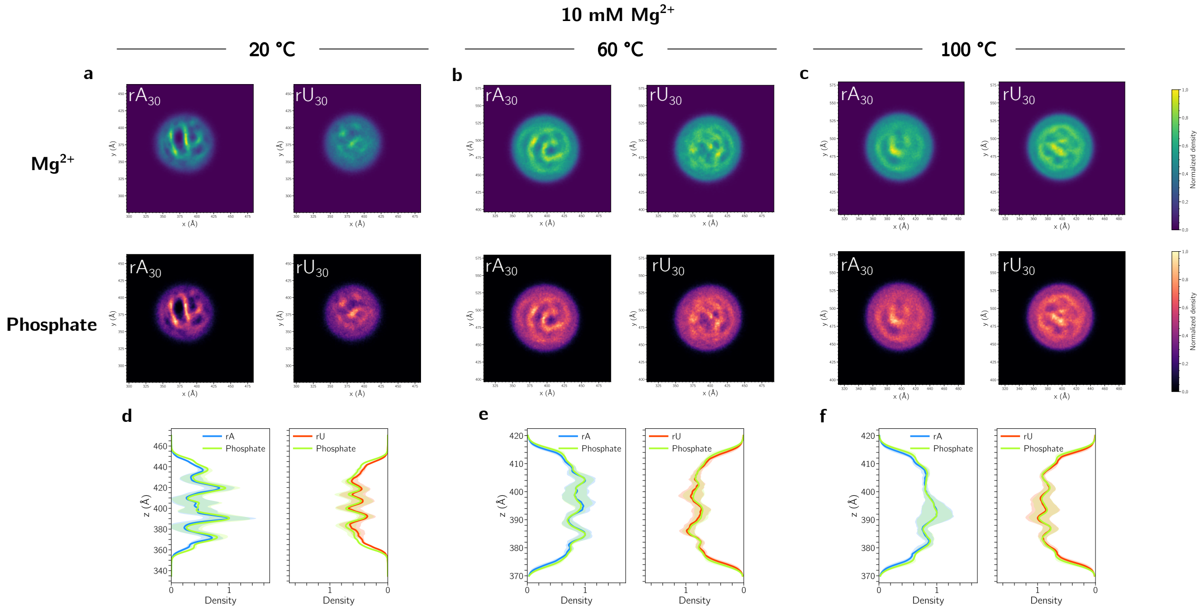

Figure S30: Same as Fig. S28, but in 10 mM $Mg^{2+}$.

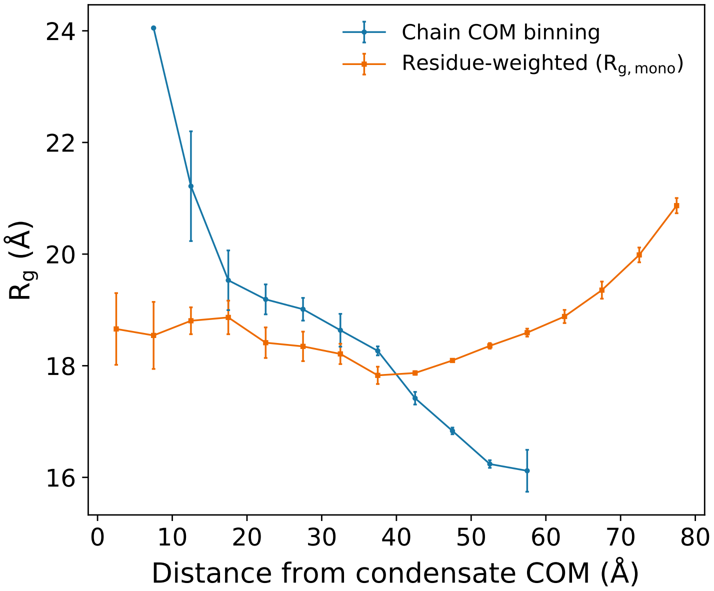

Figure S31: Comparison between two methods of calculating radial conformational profiles. Binning by the COM of the chain show that the chain becomes more compact towards the interface (in blue), whereas residue-weighted method shows non-monotonic behavior (in orange).

Movie S1: Heterogeneous distribution of RNA and $Mg^{2+}$ in $rA_{30}$ condensates.

Movie S2: Difference of internal organization between $rA_{30}$ and $rU_{30}$ condensates.
